# Pitching single-focus confocal data analysis one photon at a time with Bayesian nonparametrics

**DOI:** 10.1101/749739

**Authors:** Meysam Tavakoli, Sina Jazani, Ioannis Sgouralis, Omer M. Shafraz, Sanjeevi Sivasankar, Bryan Donaphon, Marcia Levitus, Steve Pressé

## Abstract

Fluorescence time traces are used to report on dynamical properties of molecules. The basic unit of information in these traces is the arrival time of individual photons, which carry instantaneous information from the molecule, from which they are emitted, to the detector on timescales as fast as microseconds. Thus, it is theoretically possible to monitor molecular dynamics at such timescales from traces containing only a sufficient number of photon arrivals. In practice, however, traces are stochastic and in order to deduce dynamical information through traditional means–such as fluorescence correlation spectroscopy (FCS) and related techniques–they are collected and temporally autocorrelated over several minutes. So far, it has been impossible to analyze dynamical properties of molecules on timescales approaching data acquisition without collecting long traces under the strong assumption of stationarity of the process under observation or assumptions required for the analytic derivation of a correlation function. To avoid these assumptions, we would otherwise need to estimate the instantaneous number of molecules emitting photons and their positions within the confocal volume. As the number of molecules in a typical experiment is unknown, this problem demands that we abandon the conventional analysis paradigm. Here, we exploit Bayesian nonparametrics that allow us to obtain, in a principled fashion, estimates of the same quantities as FCS but from the direct analysis of traces of photon arrivals that are significantly smaller in size, or total duration, than those required by FCS.

## I. INTRODUCTION

Methods to capture static molecular structures, such as super-resolution microscopy [36, 51, 67], provide only snapshots of life in time. Yet life is dynamical and obtaining a picture of life in action–one that captures diffraction-limited biomolecules as they move, assemble into and disassemble from larger bimolecular complexes– remains an important challenge [60]. In fact, the creative insights directly leading to fluorescence correlation spectroscopy (FCS) [30, 69]–and related methods such as FCS-FRET [84, 104] and FCCS [92]–have shown that deciphering dynamical information from molecules, often biomolecules, does not demand spatial resolution or spatial localization. Rather, the key is to inhomogeneously illuminate a sample over a small volume.

As fluorescently-labeled molecules diffuse across this inhomogeneously illuminated volume, they emit photons (*i.e.*, they fluoresce) in a way that is proportional to the illumination at their respective locations [57]. Single photon detectors, often photo-multiplier tubes or avalanche photodiodes, are then used to record these photons. In principle, with the appropriate electronics, photons can be recorded within *μs*-*ms*. This suggests that information on the molecules’ motion could be drawn from the data on fast timescales that approach data acquisition, *i.e.* no more than a few *μs*-*ms*.

The fundamental quantities measured in a confocal optical setup are individual photon arrival times, from which photon inter-arrival times, *i.e.*, the intervals between adjacent photon arrivals, can be readily obtained [59]. When imaging molecules fixed in space and under homogeneous (uniform) illumination, these inter-arrival times–excluding other experimental and label artifacts such as detector noise, background photons, and label photo-physical kinetics–are independent and identically distributed and so uncorrelated with each other. However, inter-arrival times measured in conventional confocal experiments encode the number of molecules in the vicinity of the confocal volume, their diffusion dynamics, their position with respect to the confocal center in addition to an array of experiment specific artifacts such as detector characteristics and label photo-kinetics. Consequently, inter-arrival times are correlated with each other and, in principle, these correlations can be exploited to characterize the dynamics of the underlying molecular system.

Thus far, correlations in the inter-arrival times are exploited by collecting photons over long periods [85] and temporally autocorrelating the resulting fluorescence intensity measurements [15, 30, 35, 69]. For sufficiently long intensity traces, the stochasticity in the number of labeled molecules contributing photons, as well as their positions in the illuminated volume and their instantaneous photon emission rates, are averaged out. As such, the mathematical expression for the fluorescence intensity time-autocorrelation function takes a simple form that–under strong assumptions on the illuminated volume’s geometry and the molecules’ photon emission rate– can be summarized in analytic formulas that are fitted on the acquired measurements.

However, despite the elegance and simplicity of the mathematics involved in the derivation of the time-autocorrelation function [15, 30, 35, 69], a critical limitation of autocorrelative methods, including all those within the FCS framework, remains the stark timescale separation between data collection (*e.g.*, typical time between successive photon arrivals) and the timescale required to deduce a meaningful dynamical interpretation (*e.g.*, typical duration between first and last photon arrivals used); see Fig. (1). A method that takes direct advantage of single photon arrivals, without using intensity traces (*i.e.*, downsampled photon arrivals), has the potential to reveal dynamical information on timescales several orders of magnitude faster than traditional FCS analysis. As a result, rapid or non-equilibrium processes and, as such, abrupt changes in molecular chemistry, could be studied. Furthermore, provided such a method can utilize substantially shorter traces, the total duration of experiments can be shrunk and the phototoxic damage induced on biological samples can be reduced substancially [67, 70, 83, 103]. This is especially relevant for *in vivo* FCS applications [28, 80, 93, 105].

**FIG. 1.**
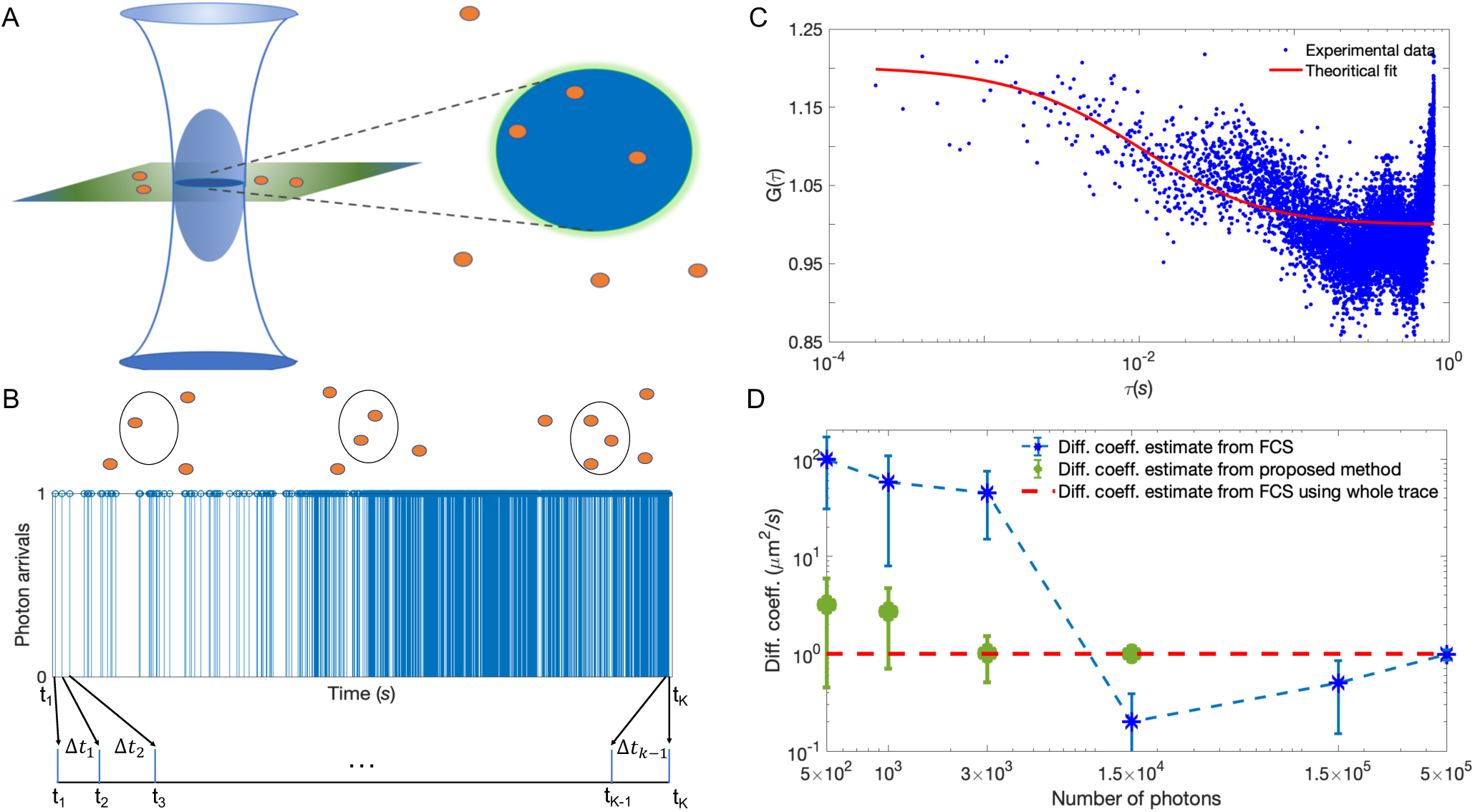
Photon arrival times can characterize dynamical properties of molecules on fast, photon-detection, timescales. (A) Schematic of an illuminated confocal volume (blue) with fluorescent molecules emitting photons based on their location within that volume. (B) Synthetic trace containing ≈ 1500 photon arrivals produced by 4 molecules diffusing at 1 *µm*^2^*/s* for a total time of 30 *ms* under background and molecule photon emission rates of 10^3^ photons/*s* and 4 × 10^4^ photons/*s*, respectively. (C) Autocorrelation curve, *G*(*τ*), of the trace in (B), binned at 100 *µs*. On account of the limited data available in the trace, any reasonable fit is impossible. Normally, in FCS analysis, much longer traces are used to generate smoother *G*(*τ*) that are fitted to determine a diffusion coefficient. In Fig. A1 of the Appendix, we show that the quality of the fit does not improve considerably by fitting to a semi-logarithmic curve. (D) Comparison between diffusion coefficient estimates using our proposed method (detailed later) and FCS as a function of the number of photon arrivals in the analyzed trace. Since by 1.5 × 10^4^ photon arrivals our method has converged, we avoid analyzing larger traces.

Previously proposed methods to analyze single photon measurements [2, 4, 40, 41, 45, 75, 76, 82, 108] make assumptions that render them inappropriate for imaging molecules moving through inhomogeneously illuminated volumes [40]. For example, for the analysis of single molecule fluorescence resonance energy transfer (FRET), existing methods assume that the photon inter-arrival times reflect only biomolecular conformational transitions [40–42] but not diffusive motion of the entire biomolecule [26, 27, 39, 41], and so are appropriate only for experiments on immobilized molecules. Along the same lines, existing methods combine FRET with FCS [90] to quantify *ns* dynamics; however, they do not directly exploit single photon measurements. Rather, they operate on downsampled measurements, achieved through binning, similar to traditional FCS, and therefore inherit the same limitations and drawbacks.

To be able to use single photon arrival times to estimate the diffusion coefficient of labeled molecules in a confocal experiment, as in most biological applications, we must be able to determine the particular number of molecules responsible for the observed photon arrival time trace. Otherwise, naively, many molecules with low diffusion coefficients emitting photons at the periphery of the illuminated confocal volume could be mistaken for fewer molecules with higher diffusion coefficients in the center region which is most illuminated. As we illustrate in Fig. (2), misidentifying the number of molecules, or incorrectly assessing their positions, may give rise to incorrect diffusion coefficient estimates.

**FIG. 2.**
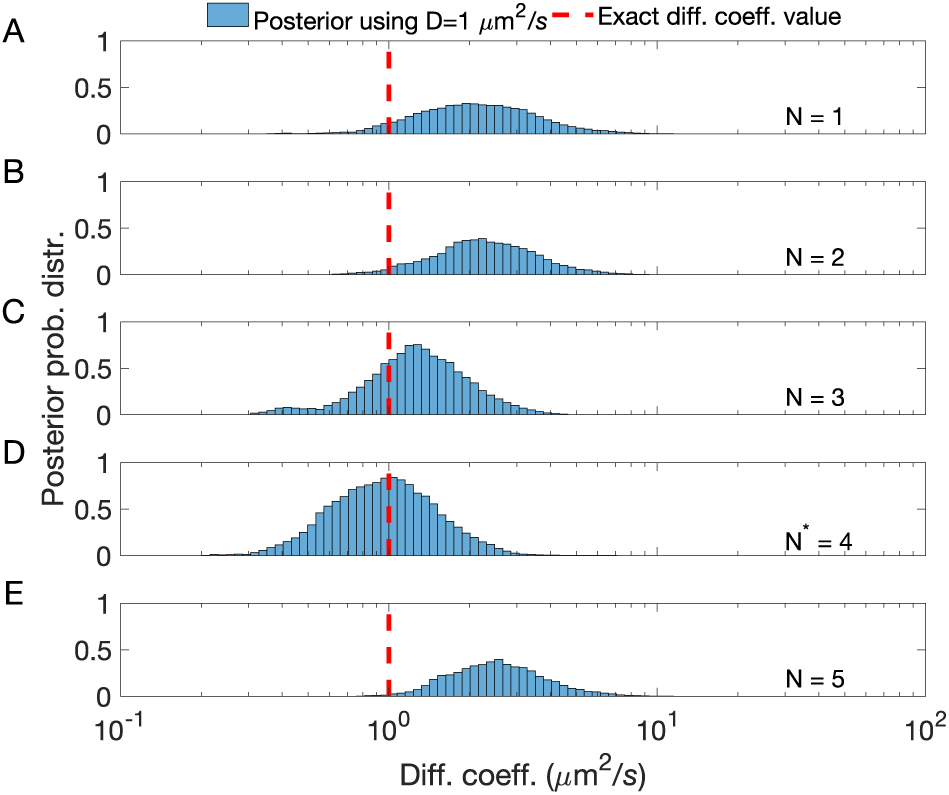
Estimates of diffusion coefficients from photon arrival traces strongly depend on the number of molecules assumed to be contributing to the trace. The trace analyzed contained ≈ 1800 photon arrivals produced by 4 molecules diffusing at 1 *μm*^2^*/s* for a total time of 30 *ms* under background and molecule photon emission rates of 10^3^ photons/*s* and 4 × 10^4^ photons/*s*, respectively. To estimate *D parametrically*, we assumed a fixed number of molecules, *N* = 1 (A); *N* = 2 (B); *N* = 3 (C); *N* = 4 (D); and *N* = 5 (E). The correct estimate in (D)–and the mismatch in all others–underscores why it is critical to estimate the number of molecules contributing to the trace to deduce quantities such as diffusion coefficients from single photon arrivals.

More concretely, to obtain quantitative estimates of the diffusion coefficient, we need to formulate a likelihood [9, 37, 107]. In turn, to formulate a likelihood for photon arrival data demands that we know the number of molecules contributing photons as well as their locations across time. As the number of molecules instantaneously located within the confocal volume is unknown, all reasonable possibilities need to be considered and rank-ordered using expensive pre- or post-processing model selection heuristics [60, 100]. This has not been achieved yet, in part, because of the prohibitive computational cost it entails. Analyzing single photon arrivals from a confocal setup to derive dynamical information therefore demands fundamentally new tools.

The conceptually novel framework that we propose in this study can winnow down infinite possibilities (*i.e.*, infinite populations of molecules potentially contributing photons) to a finite, computationally manageable, number in a mathematically exact manner. Such a framework avoids compromising temporal resolution, as it requires no intensity trace to be formed (*i.e.*, no downsampling), and allows us to directly deduce dynamical quantities, such as diffusion coefficients, efficiently from raw single photon arrivals. The underlying theory, Bayesian non-parametrics (BNPs) [34], is a powerful set of tools still under active development and largely unknown to the Physical Sciences [19, 49, 53, 60, 79, 95–98, 100].

Mathematical devices within BNPs, such as the beta-Bernoulli process [1, 16, 77], allow us to place priors not only on parameters themselves, as traditional parametric Bayesian methods, but also on distributions over an infinite number of candidate models to which parameters are associated [50]. Concretely, for the case of our single photon time traces, BNPs and in particular beta-Bernoulli processes can be used to assign posterior probabilities over an array of quantities including all possible number of molecules responsible for producing the data and their associated locations at each photon arrival time. With these devices, as we describe herewith, we turn the otherwise difficult problem of model-selection–that is, determining how many molecules contribute photons–into a parameter estimation problem that remains computationally tractable [1, 16, 77].

## II. MATERIALS AND METHODS

Here, we describe the mathematical formulation of our BNPs method for the analysis of confocal single photon data. We begin with the overall input which consists of photon inter-arrival times, **Δ*t*** = (Δ*t*_1_, Δ*t*_2_, …, Δ*t*_*K*−1_) where Δ*t*_*k*_ represents the time interval between adjacent observations of photons, which occur at times *t*_*k*_ with *k* = 1, …, *K*. We also use as input the illuminated confocal volume’s shape and background photon emission rate which we can determine separately through calibration [13].

To derive estimates for the diffusion coefficient from **Δ*t***, we need to determine intermediate quantities which include: i) photon emission rates of molecular labels; and, most importantly, ii) the unknown number of molecules contributing photons to the trace **Δ*t***, as well as their location with respect to the center of the confocal volume.

A graphical summary of our formulation is shown in Fig. (3). Below, we explain briefly each step involved. More details, and an implementation of the whole method, are available in the Appendix. In addition, source code and a GUI version of our implementation are provided through the Supplementary Materials.

**FIG. 3.**
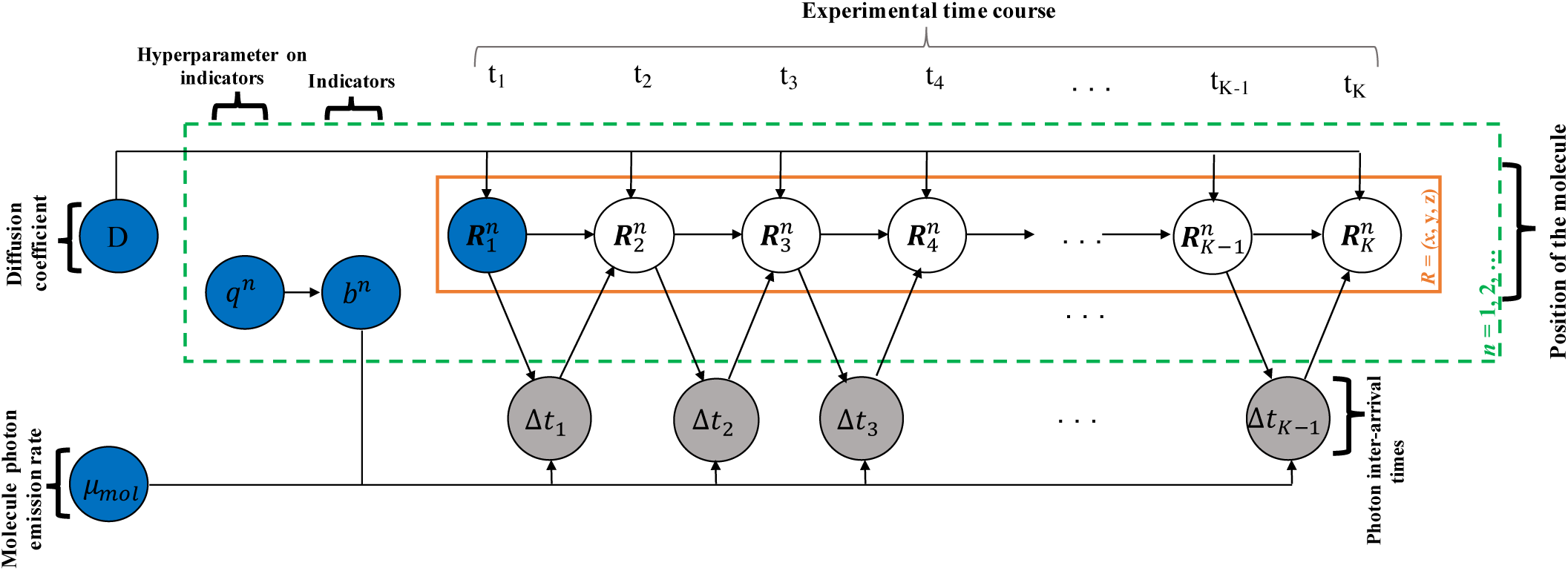
BNP formulation used for the analysis of photon arrival traces. Molecules, indexed *n* = 1, 2, …, evolve over the experimental time course which is indexed by *k* = 1, 2, …, *K*. Here, 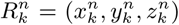 indicates the location of molecule *n* at time *t*_*k*_. During the experiment, only a single observation (inter-arrival time) Δ*t*_*k*_ is recorded, thereby combining photon emissions from every molecule and the background. The diffusion coefficient *D* determines the evolution of the molecular positions which influence the photon emission rates and eventually the recorded Δ*t*_*k*_. The indicator variables *b*^*n*^ are introduced to infer the unknown molecule population size. In the graphical model, the measured data are highlighted by grey shaded circles and the model variables, which require priors, are designated by blue circles.

### A. Model Formulation

We begin with the distribution according to which the *k*^*th*^ observation, Δ*t*_*k*_, is derived

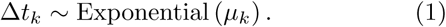

Accordingly, Δ*t*_*k*_ follows an exponential probability distribution [42, 81] with rate *μ*_*k*_. In fact, the rate *μ*_*k*_ gathers the photon emission rates of all molecules which depend on their respective locations relative to the confocal center (see below) [13]. In addition to the molecule photon emissions rates, *μ*_*k*_ also includes background photons

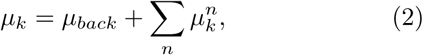

where 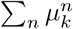 is the sum over photon emission rates 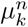 gathered from the individual molecules, that we index with *n* = 1, 2, …, and *μ*_*back*_ is the background photon emission rate. In our formulation, 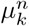 and *μ*_*back*_ are the emission rates of photons that reach our detectors which, due to optical and detector limitations, are typically lower than the rates of actual photon emissions [71, 72].

Next, we incorporate the dependency of the emission rate 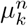 on location [5, 22, 111] with other effects such as camera pinhole shape and size, the laser intensity, laser wavelength, and quantum yield [13] into a characteristic point spread function (PSF). To be more precise, a PSF characterizes the optical response of an imaging system [12, 59, 114]. Although this term is mostly used for wide-field microscopes to describe the emission PSF, here, we follow the FCS literature, and use it to describe the confocal microscope, *i.e.*, both emission and detection PSFs. Consistent with FCS [15, 30, 35, 69], we assume a 3D Gaussian geometry [57]

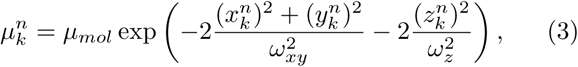

where 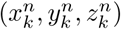 is the position of the *n*^*th*^ molecule at time *t*_*k*_ and the parameter *μ*_*mol*_ indicates the brightness of a single molecule. This is the rate of detected photon emissions achieved when the molecule is at the center of the confocal volume where illumination is highest.

Finally, for a molecule diffusing along one direction, the probability distribution *p*(*x, t*) of its position *x* at time *t* satisfies the diffusion equation [6, 21, 55]

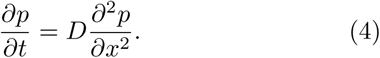

To solve this equation, we assume that the molecule is located at *x*_*k*−1_ at time *t*_*k*−1_ and we obtain

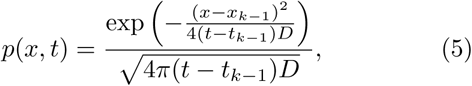

which is the probability density of a normal random variable with mean *x*_*k*−1_ and variance 2(*t* − *t*_*k*−1_)*D*. Therefore, at time *t* = *t*_*k*_, we write

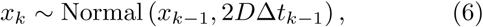

where Δ*t*_*k*−1_ = *t*_*k*_ − *t*_*k*−1_ and *D* is the molecule’s diffusion coefficient. Similarly, solving the diffusion equation for molecules following isotropic diffusion in free space along all three Cartesian directions, we obtain

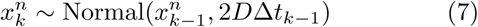

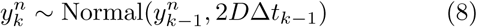

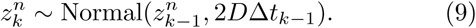

### B. Model Inference

All quantities which we need to infer–such as the diffusion coefficient, *D*, locations of molecules through time, 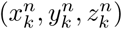 and the molecule photon emission rate *μ*_*mol*_– are formulated as model variables. We estimate these variables within the Bayesian paradigm [37, 60, 100]. The model parameters such as *D* and *μ*_*mol*_ require priors. Additionally, we have to consider priors on the initial molecule locations, *i.e.*, at the time of the very first photon arrival, 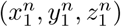. Options for these priors are straightforward and, for computational convenience, we adopt the distributions described in the Appendix.

Meanwhile, before we proceed any further with our BNPs formulation, we need to revise eq. (3) as follows

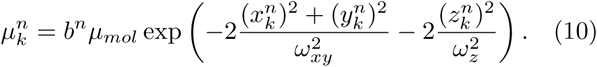

The variables *b*^*n*^, defined for each model molecule, take only values 1 or 0. Specifically, we have *b*^*n*^ = 0 when the *n*^*th*^ model molecule *does not* contribute photons to the measurements as in this case the molecule is *decoupled* from the overall photon emission rate *μ*_*k*_. This indicator variable allows us to operate on an arbitrarily large population of model molecules; technically, an infinite population. The ability to recruit, from a potentially infinite pool of model molecules, the precise number that contributes to the measured trace **Δ*t*** is the chief reason we abandon the parametric Bayesian paradigm and adopt BNPs. After introducing the indicators *b*^*n*^, we can estimate the number of molecules that contribute photons, *i.e.*, those molecules where *b*^*n*^ = 1, simultaneously with the remaining of the parameters simply by having each *b*^*n*^ as a separate parameter and estimating its value.

To estimate *b*^*n*^, we consider a Bernoulli prior with a beta hyper-prior

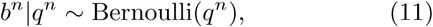

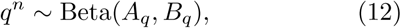

where *A*_*q*_ and *B*_*q*_ are (hyper-hyper-)parameters specifically chosen to allow for *n* → ∞. In this limit, eqs. (11) and (12) can be combined resulting in a beta-Bernoulli process [1, 16, 77]; see Appendix for more details.

With the specified priors, we can now form a joint posterior probability 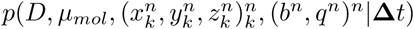 including all unknown variables which we seek to determine. Nevertheless, the nonlinear dependence of the PSF on the molecules’ positions 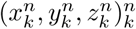 and the non-parametric prior on the indicators (*b*^*n*^)^*n*^ exclude a closed form for our posterior. For this reason, we develop a Markov Chain Monte Carlo scheme [37, 58, 86] that exploits results from the theory of Computational Statistics and Non-linear filtering to generate pseudo-random samples from this posterior that we use in obtaining our estimates [37, 86]. A technical description of this scheme can be found in the Appendix and a ready-to-use implementation is available through the Supplementary Materials.

### C. Data Acquisition

#### 1. Acquisition of Synthetic Data for Figs. (4)-(7)

We acquire the synthetic data shown in the Results section by computer simulations [8, 33, 44, 47, 52] that represent Brownian motion of point molecules moving through a typical illuminated confocal volume. We provide finer details and complete parameter choices in the Appendix.

**FIG. 4.**
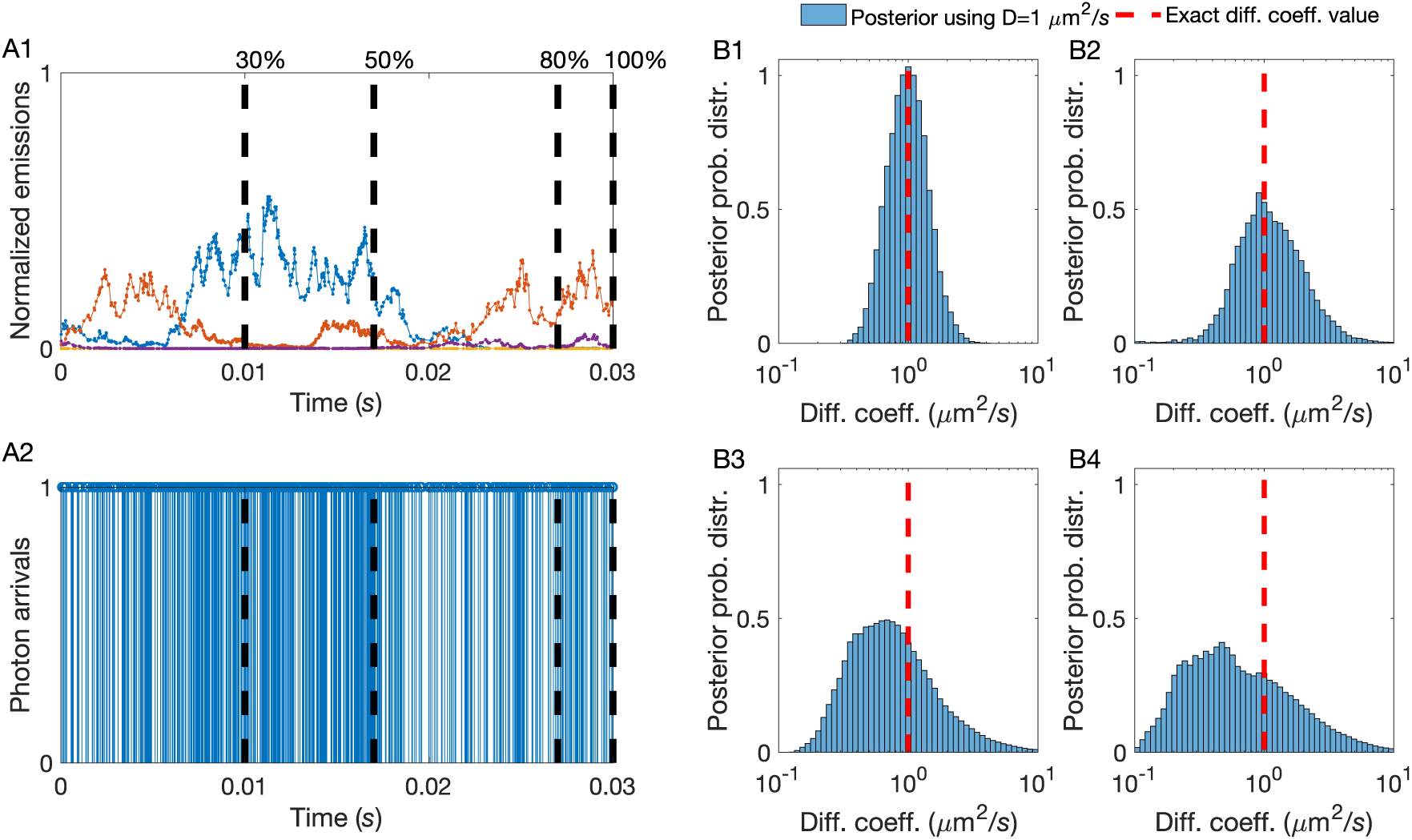
A higher number of total photon arrivals provide more photons per unit time and sharper diffusion coefficient estimates. (A1) Instantaneous molecule photon emission rates 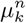, normalized by *μ*_*mol*_. (A2) Photon arrival trace resulting from combining photon emissions from every molecule and the background. This synthetic trace contains ≈ 2000 photon arrivals produced by 4 molecules diffusing at 1 *μm /s* for a total time of 30 *ms* under background and molecule photon emission rates of 10^3^ photons/*s* and 4 × 10^4^ photons/*s*, respectively. The dashed lines show the initial 30%, 50%, 80%, and 100% portions of the original trace containing ≈ 600, ≈ 1000, ≈ 1600, ≈ 2000 photon arrivals, respectively. (B1-B4) Posterior probability distributions drawn from traces with differing length (shown in (A2)). As expected, for the longer traces, the peak of the posterior matches with the exact value of *D* (dashed line). Gradually, as we decrease the total number of photon arrivals analyzed, the estimation becomes less reliable.

**FIG. 5.**
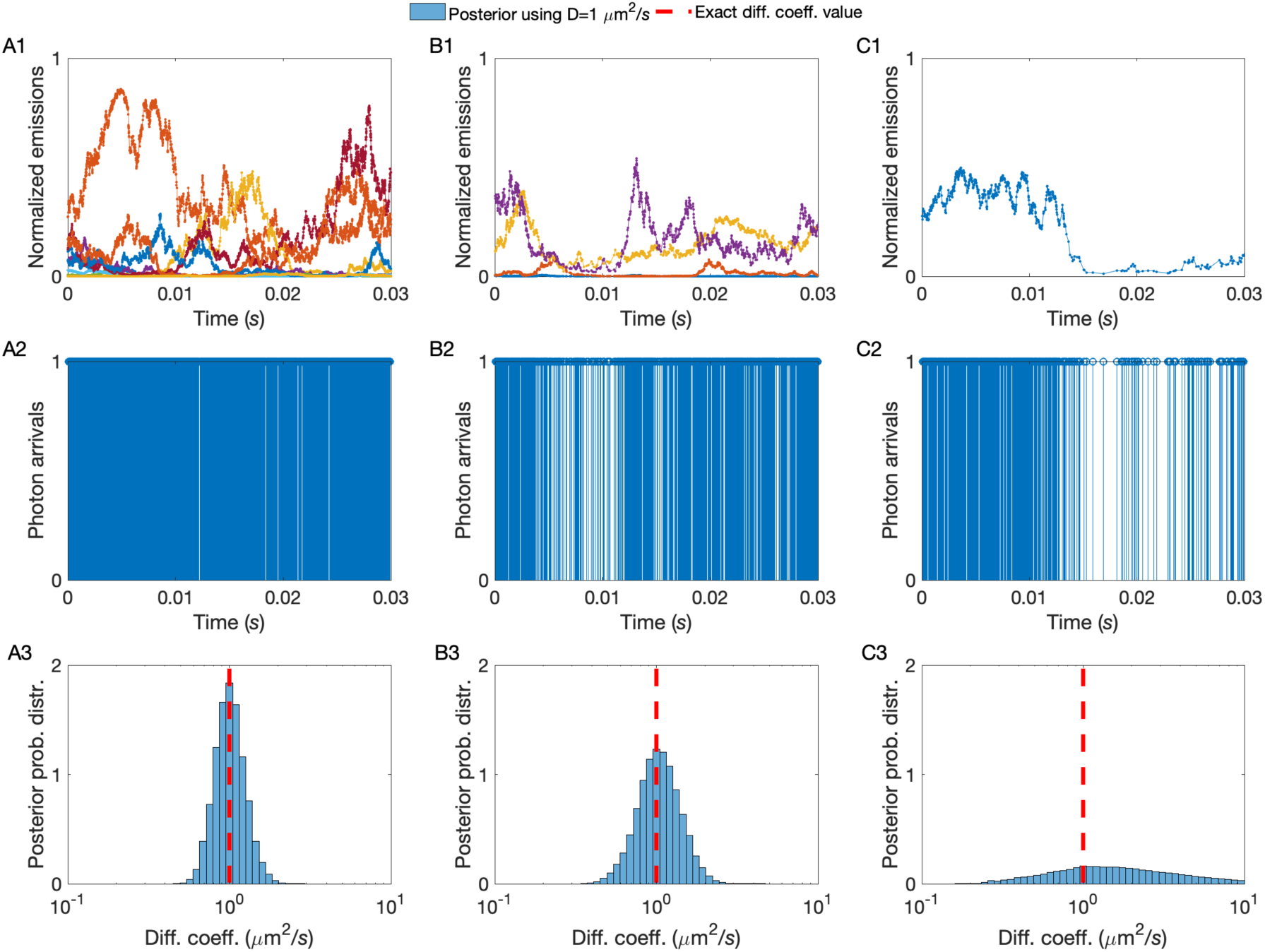
A higher molecular concentration provides more photons per unit time and sharper diffusion coefficient estimates. (A1, B1, C1) Instantaneous molecule photon emission rates 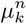, normalized by *μ*_*mol*_. (A2, B2, C2) Photon arrival traces resulting from combining photon emissions from every molecule and the background. These are produced by 10 molecules containing ≈ 3000 photon arrivals (A2), 4 molecules containing ≈ 2000 photon arrivals (B2), and 1 molecules containing ≈ 1000 photon arrivals (C2), diffusing at 1 *μm /s* for a total time of 30 *ms* under background and molecule photon emission rates of 10^3^ photons/*s* and 4 10^4^ photons/*s*, respectively. (A3, B3, C3) Posterior probability distributions drawn from traces with differing number of molecules (shown in (A2, B2, C2)). As expected, for the traces with higher number of molecules, the peak of the posterior matches with the exact value of *D* (dashed line). Gradually, as we decrease the total number of molecules the estimation becomes less reliable.

**FIG. 6.**
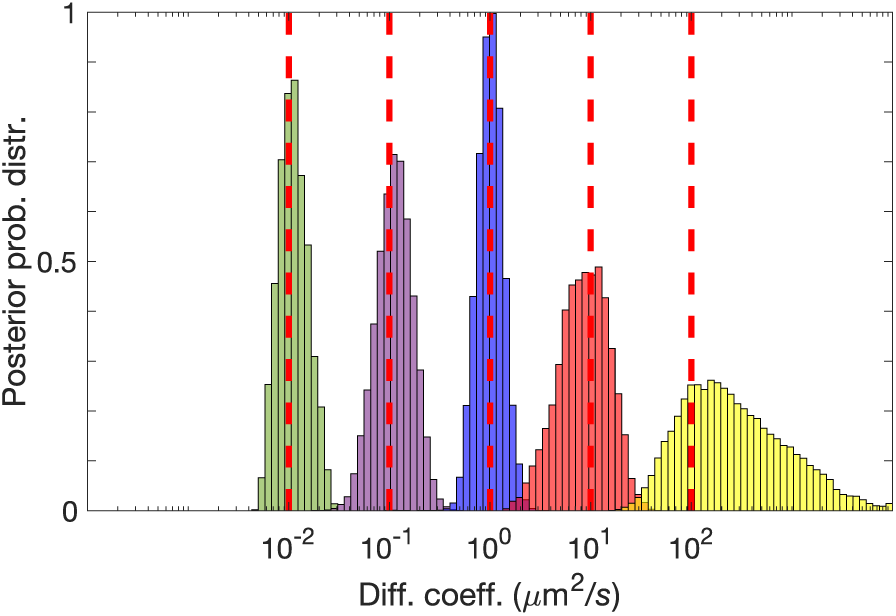
A lower diffusion coefficient provides more photons per unit time and sharper diffusion coefficient estimates. Posterior probability distributions drawn from traces containing ≈ 2000 photon arrivals produced by 4 molecules diffusing at *D* = 0.01, 0.1, 1, 10 *μm*^2^*/s* for a total time of 30 *ms* under background and molecule photon emission rates of 10^3^ photons/*s* and 4 × 10^4^ photons/*s*, respectively. For molecules diffusing at *D* = 100 *μm*^2^*/s*, under similar conditions, we used a trace containing ≈ 3000 photons for a total time of 50 *ms*, since we needed a longer trace to gather sufficient information for drawing a posterior.

**FIG. 7.**
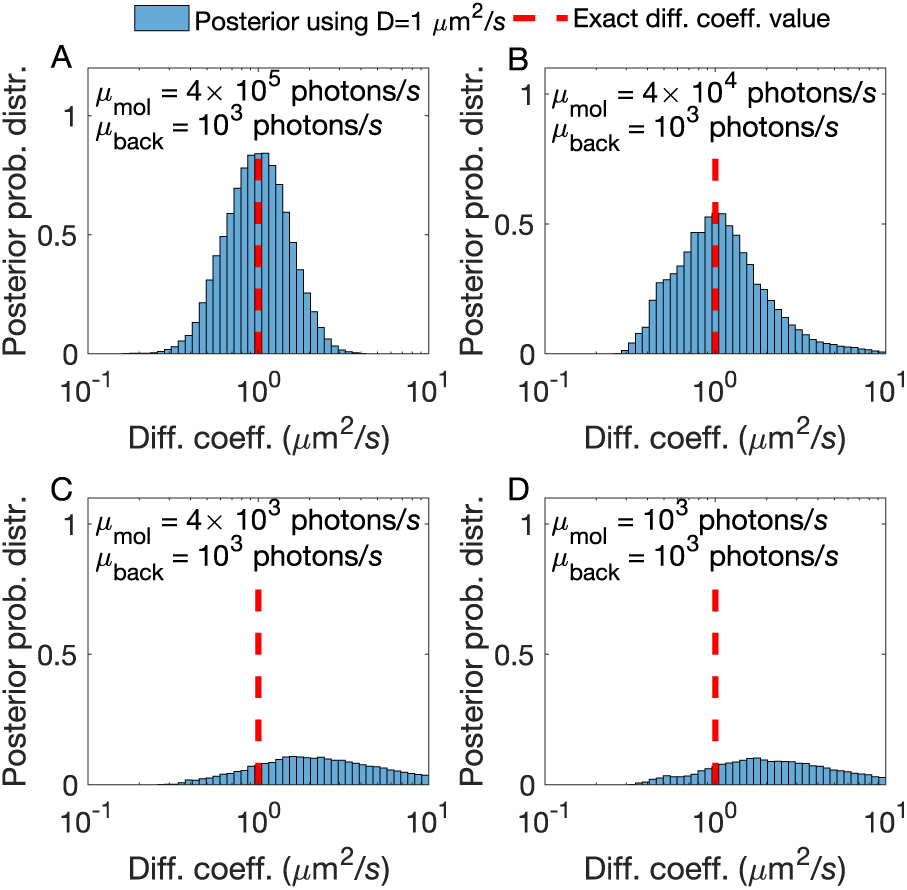
A higher molecule photon emission rate provides more photons per unit time and sharper diffusion coefficient estimates. (A, B, C, D) Posterior probability distributions drawn from traces produced by 4 molecules diffusing at 1 *μm*^2^*/s* for a total time of 30 *ms* under background photon emission rate of 10^3^ photons/*s* and molecule photon emission rates 4 × 10^5^, 4 × 10^4^, 4 × 10^3^, 10^3^ photons/*s*, respectively. As expected, under higher molecule photon emission rates, the peak of the posterior matches sharply with the exact value of *D* (dashed line). Gradually, as we decrease the molecule photon emission rate, the estimation becomes less reliable.

#### 2. Acquisition of Experiment data for Figs. (8)-(12)

For these experiments we used Cy3 fluoresent dyes. Solutions were made by suspending Cy3 dye in glycerol/buffer (pH 7.5, 10 mM Tris-HCl, 100 mM NaCl and 10 mM KCl, 2.5 mM *CaCl*_2_) at various v/v, to a final concentration of either 100 pM or 1 nM. The solution was placed in a glass-bottomed fluid-cell, assembled on a custom designed confocal microscope [63] and a 532 nm laser beam was focused to a diffraction-limited spot on the glass coverslip of the fluid-cell using a 60x, 1.42 N.A., oil-immersion objective (Olympus). In our setup, the laser beam is focused at the glass-water/glycerol interface and the beam is refocused by visual inspection at the beginning of every measurement. Emitted fluorescence was collected from the same objective and focused onto a Single Photon Avalanche Diode (SPAD, Micro Photon Devices) with a maximum count rate of 11.8 Mc/s. A bandpass filter placed in front of the detector blocked all back-scattered excitation light and relayed only fluorescence from Cy3. Individual photon arrivals on the detector triggered TTL pulses and were both timestamped and registered at 80 MHz. This was achieved using a field programmable gate array (FPGA, NI Instruments) and custom LabVIEW software [89].

**FIG. 8.**
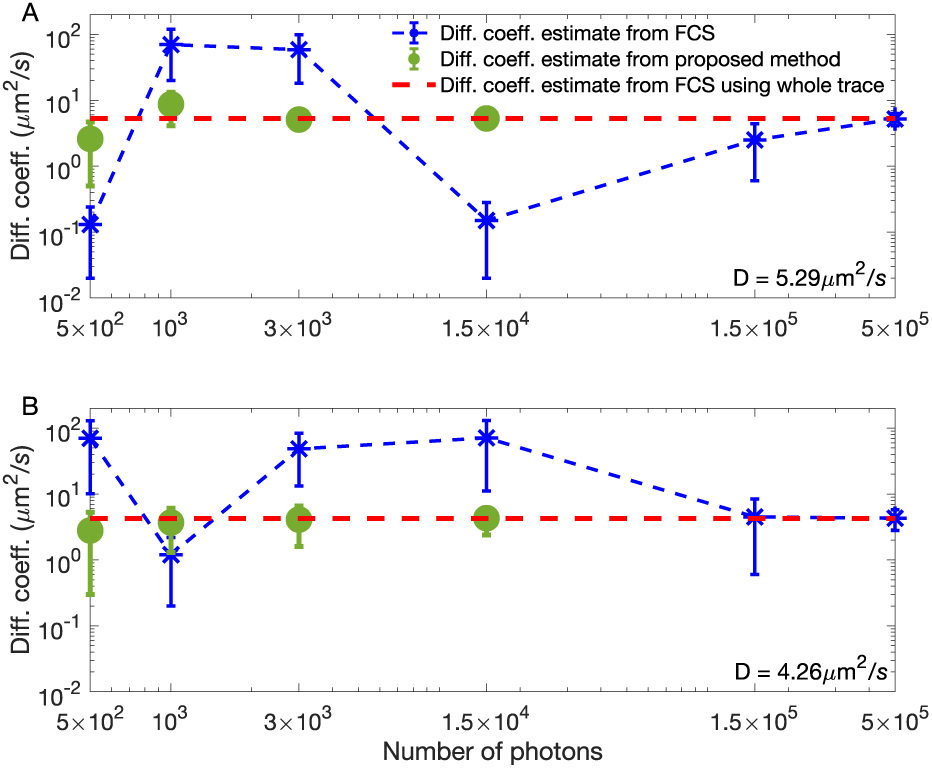
Higher molecular concentrations in experimental traces provide more photons per unit time resulting in sharper diffusion coefficient estimates. Estimates shown are drawn from experimental traces with a low (100 pM) (A) and high (1 nM) (B) concentration of Cy3 dye molecules and 75% glycerol at a fixed laser power of 100 *μW*. Similarly to Fig. (1), we compare our method’s diffusion coefficient estimate (circle green dots) to FCS (blue asterisk) as a function of the number of photons used in the analysis. Since by 1.5 × 10^4^ photon arrivals our method has converged, we avoid analyzing larger traces. The red dash line is the FCS estimate produced from the entire 5 min trace containing ≈ 3 × 10^6^ photon arrivals.

**FIG. 9.**
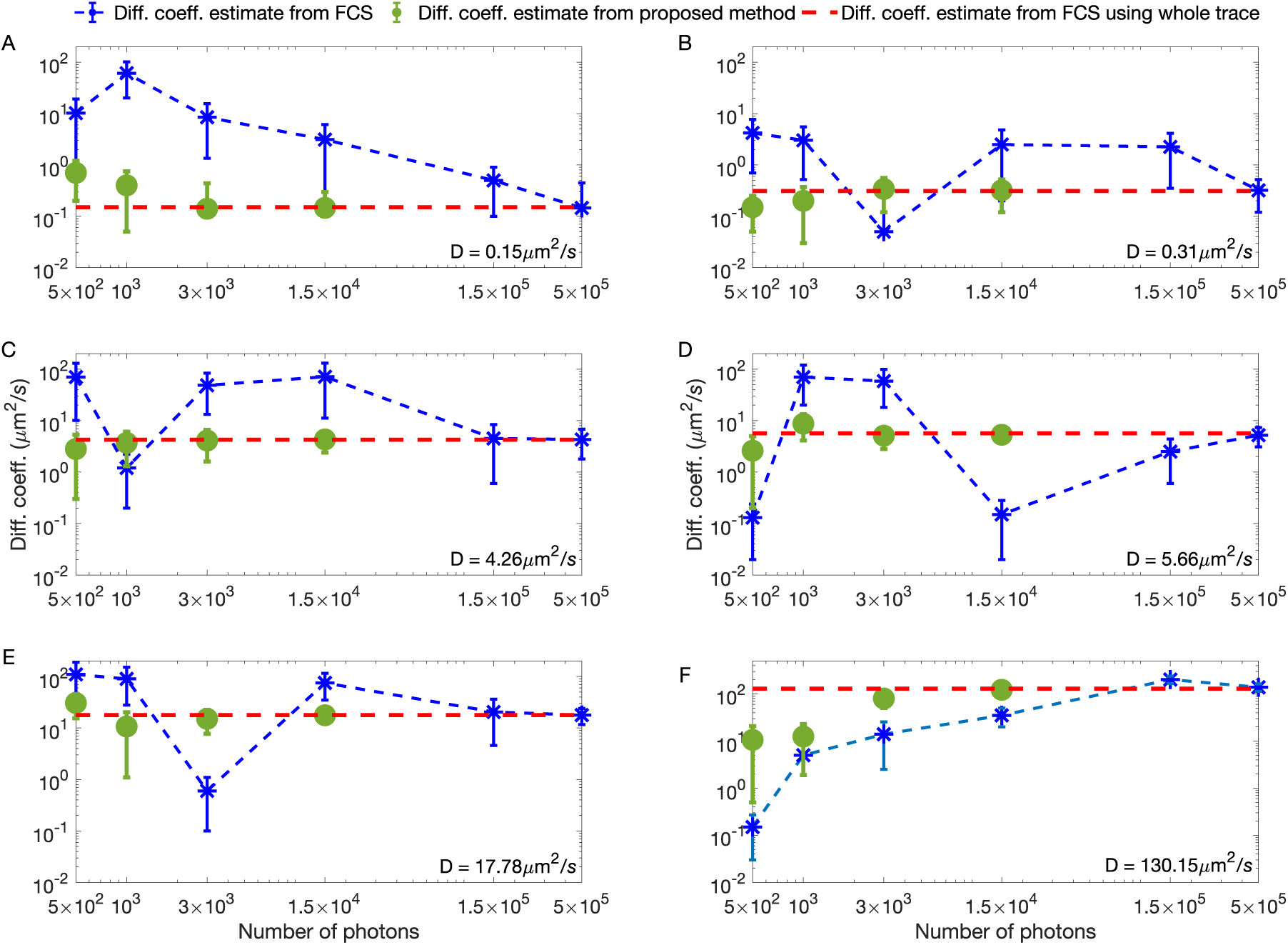
Lower diffusion coefficients in experimental traces provide more photons per unit time and sharper diffusion coefficient estimates. Estimates shown are drawn from experimental traces with 99% glycerol (A), 94% glycerol (B), 75% glycerol (C), 67% glycerol (D), 50% glycerol (E), and 0% glycerol (F) with fixed concentration 1 nM of Cy3 dye molecules and laser power of 100 *μW*. Similarly to Fig. (1), we compare our method’s diffusion coefficient estimate (circle green dots) to FCS (blue asterisk) as a function of the number of photons used in the analysis. Since by 1.5 × 10^4^ photon arrivals our method has converged, we avoid analyzing larger traces. The red dash line is the FCS estimate produced from the entire 5 min trace containing ≈ 3 × 10^6^ photon arrivals.

**FIG. 10.**
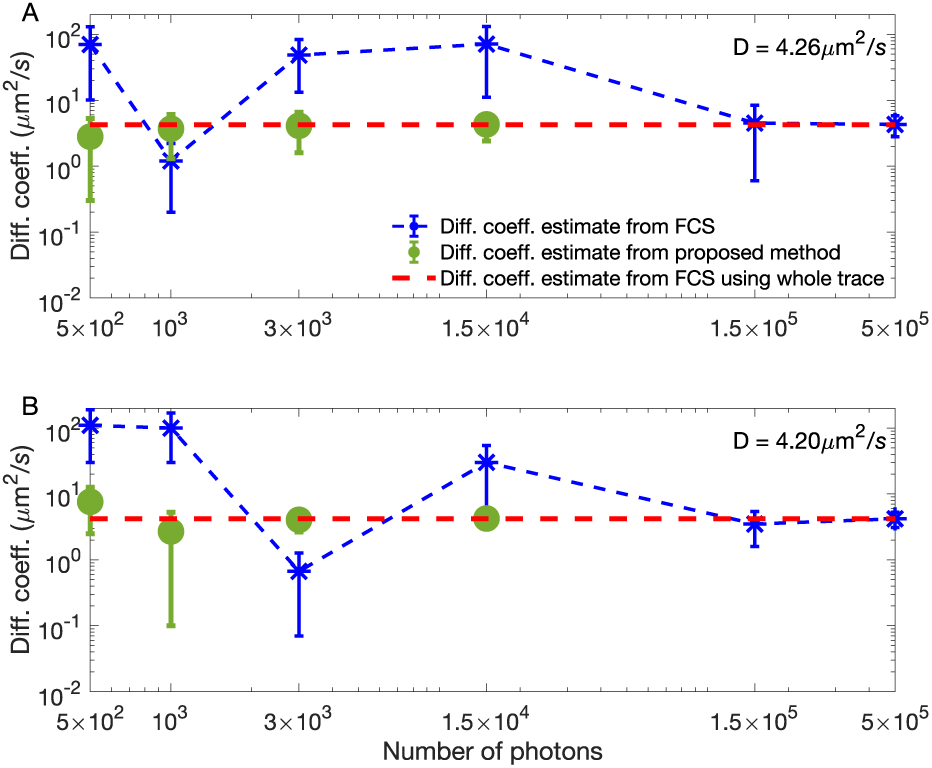
Higher laser powers in experimental traces provide more photons per unit time and sharper diffusion coefficient estimates. Estimates shown are drawn from experimental traces with high (100 *μW*) (A) and low (25 *μW*) (B) laser power with fixed concentration 1 nM of Cy3 dye molecules and 75% glycerol. Similarly to Fig. (1), we compare our method’s diffusion coefficient estimate (circle green dots) to FCS (blue asterisk) as a function of the number of photons used in the analysis. Since by 1.5 × 10^4^ photon arrivals our method has converged, we avoid analyzing larger traces. The red dash line is the FCS estimate produced from the entire 5 min trace containing ≈ 3 × 10^6^ photon arrivals.

**FIG. 11.**
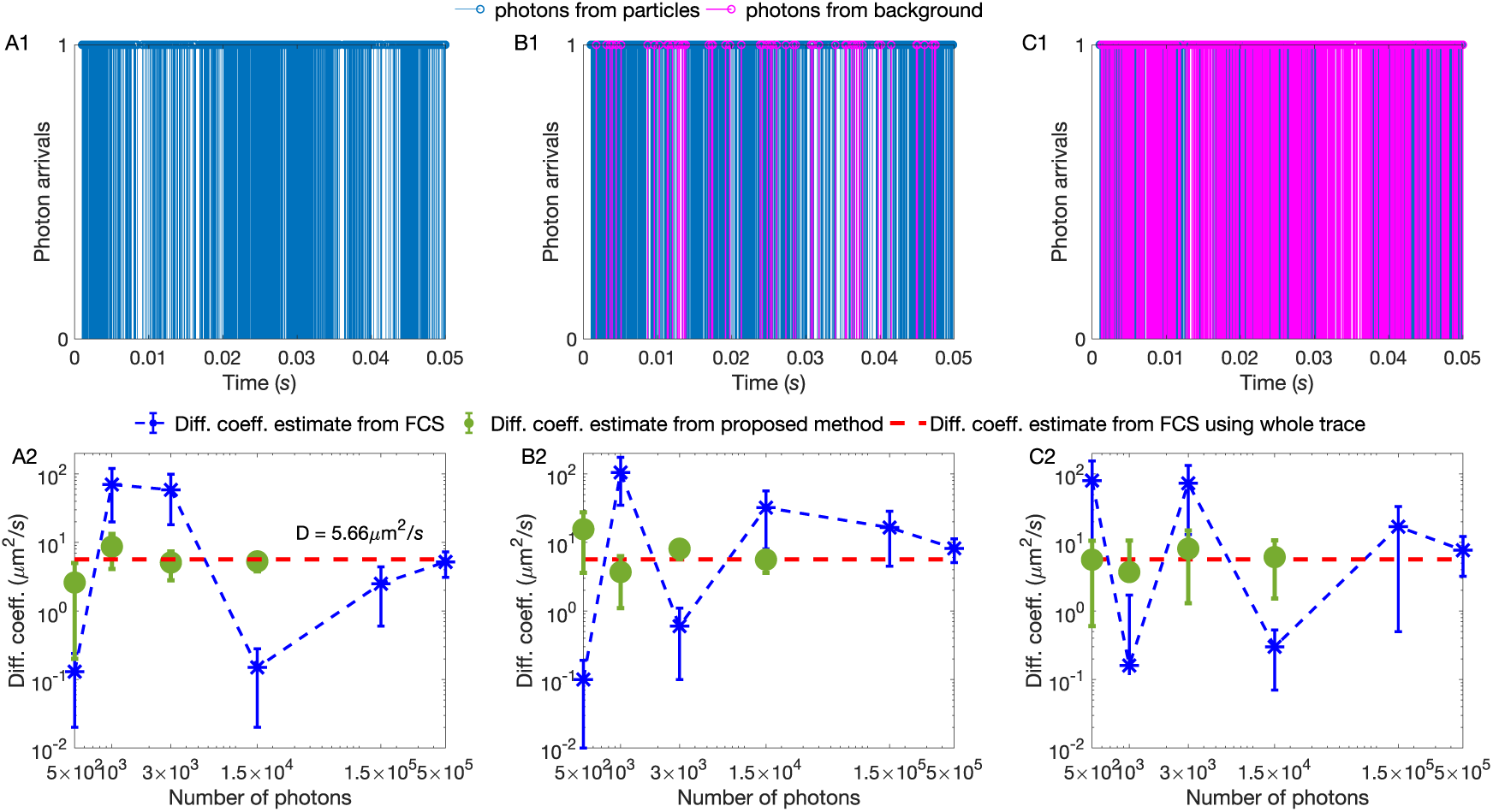
Background photon emission rates are artificially added to experimental traces yielding challenging imaging conditions and broader diffusion coefficient estimates. Experimental traces with fixed concentration 1 nM of Cy3 dye molecules and 67% glycerol and fixed laser power 100 *μW*. The same total number of photons analyzed under differing (artificially increased) background photon emission rates (0 (A1), 500 (B1), 1000 (C1) photons/*s*). (A2, B2, C2) Similarly to Fig. (1), we compare our method’s diffusion coefficient estimate (green dots) to FCS (blue asterisk) as a function of the number of photons used in the analysis. Since by 1.5 × 10^4^ photon arrivals our method has converged, we avoid analyzing larger traces. The red dash line is the FCS estimate obtained from the entire, 5 min, trace containing ≈ 3 × 10^6^ photon arrivals.

**FIG. 12.**
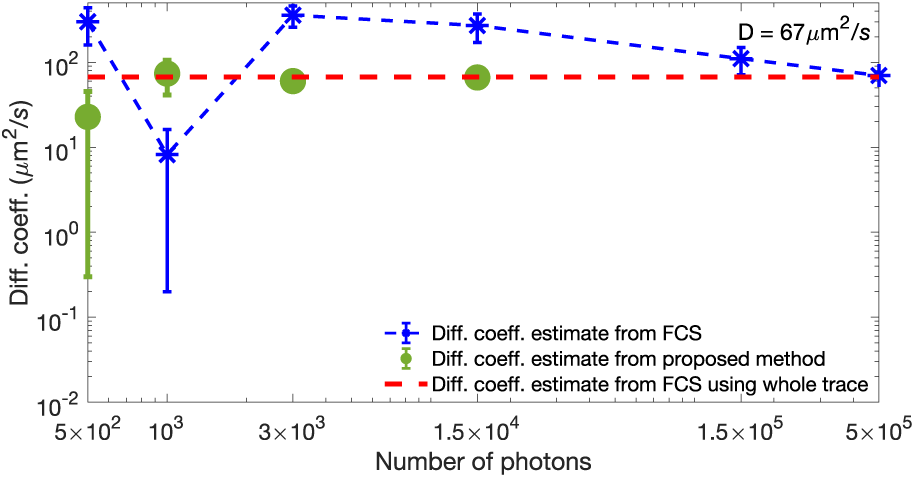
Diffusion coefficient estimates of labeled protein. Estimates shown are drawn from experimental traces with fixed concentration 1 nM of Cy3-labeled streptavidin molecules and laser power 100 *μW*. Similarly to Fig. (1), we compare our method’s diffusion coefficient estimate (green dots) to FCS (blue asterisk) as a function of the number of photons used in the analysis. Since by 1.5 × 10^4^ photon arrivals our method has converged, we avoid analyzing larger traces. The red dash line is the FCS estimate obtained from the entire, 5 min, trace containing ≈ 3 × 10^6^ photon arrivals.

#### 3. Acquisition of Experimental Data for Fig. (13)

For these experiments we used 5-TAMRA fluorescent dyes. The excitation source was a supercontinuum fiber laser Fianium WhiteLase SC480 (NKT Photonics, Birkerod, Denmark) operating at a repetition rate of 40 MHz. The excitation wavelength (550 nm) was selected by an acousto-optic tunable filter (AOTF), and the exiting beam was collimated and expanded by approximately a factor of three to slightly overfill the back aperture of the objective lens. The light was reflected into the objective lens (Zeiss EC Plan-Neofluar 100x oil, 1.3 NA pol M27, Thornwood, NY, USA) by a dichroic mirror (Chroma 89016bs). The same objective was used to collect the fluorescence from the sample, and passed through a band pass filter (Chroma ET575/50m) before being focused into a position motorized pinhole wheel set at 25 *μ*m. The output of the pinhole was focused on a multimode hybrid fiber optic patch cable (M18L01, Thorlabs, NJ, USA) which was coupled to a single-photon avalanche diode (SPCM AQRH-14, Excelitas Technologies, Quebec, Canada). The detected photons were recorded by a TimeHarp 200 time-correlated single photon counting board (PicoQuant, Berlin, Germany) operating in T3 mode. The sample (≈50 *μ*L) was contained in a perfusion chamber gasket (CoverWell) adhered on a glass coverslip. The sample was 20 nM 5-Carboxytetramethylrhodamine (5-TAMRA, purchased from Sigma-Aldrich, USA) dissolved in doubly distilled water at room temperature.

**FIG. 13.**
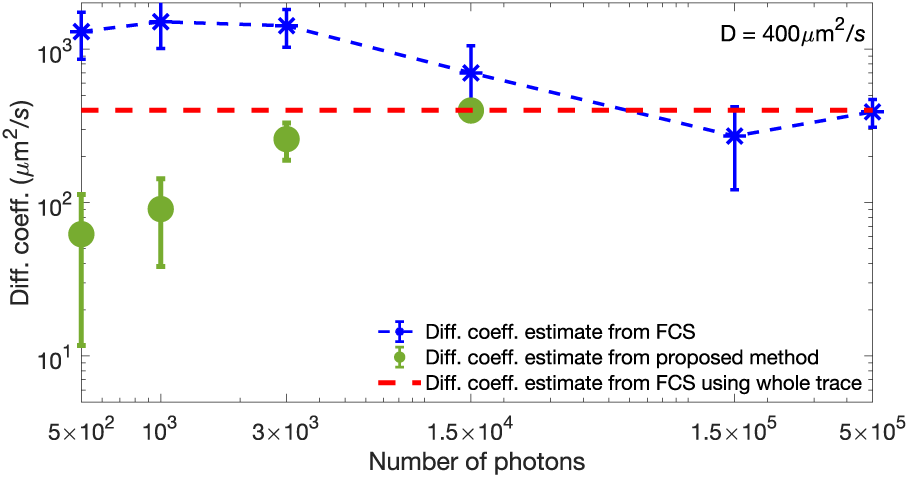
Diffusion coefficient estimates of 5-TAMRA dye. Estimates shown are drawn from experimental traces with fixed concentration 20 nM of 5-TAMRA dye molecules. Similarly to Fig. (1), we compare our method’s diffusion coefficient estimate (green dots) to FCS (blue asterisk) as a function of the number of photons used in the analysis. Since by 1.5 × 10^4^ photon arrivals our method has converged, we avoid analyzing larger traces. The red dash line is the FCS estimate obtained from the entire, 10 min, trace containing ≈ 6 × 10^4^ photon arrivals.

## III. RESULTS

Our goal is to characterize quantities that describe molecular dynamics, especially dynamics encountered in biological samples, such as diffusion coefficients, at the data-acquisition timescales of conventional single-focus confocal setups. Our input consists of: i) the measured photon inter-arrival times **Δ*t*** = (Δ*t*_1_, Δ*t*_2_, …, Δ*t*_*K*−1_); ii) the background photon emission rate; and iii) the geometry of the illuminated volume specified through a characteristic PSF.

As we explain in the Methods section, in order to estimate the molecules’ diffusion coefficient, *D*, we also estimate intermediate quantities (namely, molecule photon emission rates, molecule positions over time and the molecule numbers in the first place). These intermediate quantities demand that we use BNPs to determine quantities that *a priori* may be arbitrarily large such as the number of molecules contributing photons to our datasets **Δ*t***.

Within the Bayesian paradigm [60, 107], our estimates take the form of posterior probability distributions over the unknown quantities. These distributions combine parameter values, probabilistic relations among different parameters, as well as the associated uncertainties. According to the common statistical interpretation [37, 107], the sharper the posterior, the more conclusive (and certain) the estimate. To quantify the uncertainty, we compute a posterior variance and use the square root of this variance to construct error-bars (*i.e.*, credible intervals) [37, 107]. In Table I in the Appendix, we summarize the mean values and error bars of our analyses.

**TABLE I.** Here, we list point estimates of our analyses, which we obtain from the marginal posterior probability distributions *p*(*D*|**Δ*t***) and *p*(*μ*_*mol*_|**Δ*t***). Estimates are listed according to figure.

| | $D$ | | $\mu_{mol}$ | |
| --- | --- | --- | --- | --- |
|  | mean | std | mean | std |
| | $\mu m^2/s$ | $\mu m^2/s$ | photons/$s$ | photons/$s$ |
| Fig. (2A) | 4.54 | 4.49 | - | - |
| Fig. (2B) | 4.17 | 4.11 | - | - |
| Fig. (2C) | 1.14 | 1.12 | - | - |
| Fig. (2D) | 1.02 | 1.01 | - | - |
| Fig. (2E) | 4.75 | 4.64 | - | - |
| Fig. (4B1) | 1.03 | 0.25 | - | - |
| Fig. (4B2) | 0.95 | 0.63 | - | - |
| Fig. (4B3) | 0.75 | 0.68 | - | - |
| Fig. (4B4) | 0.45 | 0.77 | - | - |
| Fig. (5A3) | 1.01 | 0.27 | - | - |
| Fig. (5B3) | 1.09 | 0.51 | - | - |
| Fig. (5C3) | 1.65 | 1.59 | - | - |
| Fig. (6) | $1.05 \times 10^{-2}$ | $0.22 \times 10^{-2}$ | - | - |
| | $1.21 \times 10^{-1}$ | $0.34 \times 10^{-1}$ | - | - |
|  | 1.06 | 0.19 | - | - |
|  | 9.87 | 2.33 | - | - |
|  | 117.62 | 35.13 | - | - |
| Fig. (7A) | 0.99 | 0.34 | - | - |
| Fig. (7B) | 0.96 | 0.51 | - | - |
| Fig. (7C) | 2.92 | 2.68 | - | - |
| Fig. (7D) | 3.26 | 2.95 | - | - |
| Fig. (A3A3) | 10.02 | 1.17 | - | - |
| Fig. (A3B3) | 9.96 | 2.19 | - | - |
| Fig. (A4A3) | - | - | $4.11 \times 10^5$ | $1.61 \times 10^3$ |
| Fig. (A4B3) | - | - | $4.37 \times 10^4$ | $2.84 \times 10^3$ |
| Fig. (A4C3) | - | - | $1.28 \times 10^5$ | $1.25 \times 10^4$ |

Below, we validate first our method on synthetic data where the ground truth is available. For these, we use a confocal volume of typical size *ω*_*xy*_ = 0.3 *μm* and *ω*_*z*_ = 1.5 *μm* [59]. We then test our method on experimental data collected in two labs utilizing different FCS setups. For the latter cases, we demonstrate the advantages of our method by comparing our results to the results obtained from autocorrelative methods used in FCS analysis.

### A. Method Validation using Simulated Data

To demonstrate the robustness of our approach, we simulate raw single photon arrival traces under a broad range of: i) total photon arrivals, Fig. (4); ii) concentrations of labeled molecules, Fig. (5); iii) diffusion coefficients, Fig. (6); and iv) molecule photon emission rates, Fig. (7). The parameters not varied are held fixed at the following baseline values: diffusion coefficient of 1 *μm*^2^*/s* which is typical of slower *in vivo* conditions [7, 68, 88, 105], molecule photon emission rates of 4 × 10^4^ photons/*s* [62, 82], and 4 as the number of labeled molecules contributing photons. We chose 4, a small number of molecules (as opposed to a larger number of molecules), because this scenario presents the greatest analysis challenge as very few photons, and thus little data, are gathered to aid the analysis.

As illustrated in Fig. (1), a critical and recurring point throughout this section is that the traces we analyze are shorter than those that could be meaningfully analyzed using FCS. While we focus on the diffusion coefficient estimation here, we note that our framework supports more detailed parameter estimation which we provide in the Appendix.

#### 1. Total Photon Arrivals

We evaluate the robustness of our method with respect to the length of the trace (*i.e.*, the total number of photon arrivals recorded) at a fixed number of molecules, diffusion coefficient, and molecule photon emission rates. The first important finding is that, for the values of parameters selected, we need 2 orders of magnitude less data than FCS; see Fig. (1D). For instance, to obtain an estimate of the diffusion coefficient within 10% of the ground truth value, we require ≈ 10^3^ photons (directly emitted from the labeled molecule), while FCS requires ≈ 10^5^ photons. Under our simulated scenario, these correspond to traces of total duration 30 − 50 *ms* and 50 *s*, respectively. To determine our error, we chose the mean value of the diffusion coefficient’s marginal posterior, *p*(*D*|**Δ***t*), and measure the percentage difference of this mean value to the ground truth.

In general, the precise photon numbers demanded by our method and traditional FCS depend on a broad range of experimental parameter settings. This is the reason, we explore different settings in subsequent subsections as well as the Appendix.

An important overarching concept is the concept of a photon arrival as a unit of information. The more photon arrivals we have in the analyzed trace, the sharper our diffusion coefficient estimates become. This is valid, as we see in Fig. (1D) and Fig. (4), for increasing total photon arrivals. Similarly, as we see in subsequent subsections, we also collect more photons as we increase the concentration of labeled molecules (and thus the number of molecules contributing photons to the trace), increase the molecule photon emission rates of molecular labels, or decrease diffusion coefficients of molecules. In the latter case, a slower diffusion coefficients provides more time for each molecule to traverse the illuminated region, in turn, resulting in more photon arrivals.

#### 2. Molecule Concentration

To test the robustness of our method under different concentrations of labeled molecules at fixed diffusion coefficient, and molecule photon emission rates, we simulate molecules diffusing at 1 *μm*^2^/s for a total time 30 *ms* with: i) average concentrations of 10 molecules/*μm*^3^, Fig. (5A1, A2); ii) 4 molecules/*μm*^3^, Fig. (5B1, B2); and iii) 1 molecule/*μm*^3^, Fig. (5C1, C2). The molecule and background photon emission rates are taken to be 4 × 10^4^ photons/*s* and 10^3^ photons/*s* respectively, which are typical of confocal imaging [82].

Figure (5) summarizes our results and suggests that posteriors over diffusion coefficients are broader–and thus the accuracy with which we can pinpoint the diffusion coefficient drops–when the concentration of labeled molecules is lower. Intuitively, we expect this result as fewer molecules within the confocal volume provide fewer photons arrivals.

#### 3. Diffusion Coefficients

We repeat the simulations of the previous subsection to demonstrate, using synthetic data, the robustness of our method with respect to the diffusion coefficient magnitude at fixed number of molecules, and molecule photon emission rates; see Fig. (6). Intuitively, and again on the basis of the fact that photon arrivals carry information, we expect that faster moving molecules give rise to broader posterior distributions as these emit fewer photons, and thus provide less information, while they traverse the confocal volume.

#### 4. Molecule Photon Emission Rates

Figure (7) illustrates the robustness of our method with respect to the molecule photon emission rates (*i.e.*, set by the laser power used in the experimental setting and the choice of fluorescent label) by fixing the number of molecules, diffusion coefficient (1 *μm*^2^*/s*), and background emission (10^3^ photons/s). To accomplish this, we simulate increasingly dimmer molecules until the molecule signature is effectively lost in the background. As expected, dimmer molecules lead to broader posterior estimates over diffusion coefficients as these traces are associated with higher uncertainty.

### B. Estimation of Physical Parameters from Experimental Data

To evaluate our BNPs method on real data, we used experimental single photon traces collected under a broad range of conditions. That is, we used measurements from two different experimental setups and different fluorescent dyes, that are commonly used in labeling biological samples, as well as diffusing labeled proteins. Additional differences between the setups include different numerical apertures (NA), laser powers, and overall detection instrumentation as detailed in the Methods section.

Figures (8)-(11) were collected using the Cy3 dye and these results were used to benchmark the robustness of our method on dye concentration, diffusion coefficients, and laser power. Moreover, to evaluate the proposed approach beyond free dyes, in Fig. (12), we used labeled proteins, namely freely diffusing streptavidin labeled with Cy3. For Fig. (13), photon arrivals were collected using 5-TAMRA dye in order to test the robustness of our method on a different fluorophore.

#### 1. Benchmarking on Experimental Data using Cy3

We begin by verifying our method on mixtures of water and glycerol. While we only use short segments in our analysis, the collected traces are long enough (≈5 min each) to be meaningfully analyzed by traditional auto-correlative analysis used in FCS for sake of comparison. The result of the analysis of the full trace by FCS yields a diffusion coefficient that we treat as an effective ground truth. We then ask how long of a trace our method requires, as compared to FCS, in order for our diffusion coefficient estimate to converge to this ground truth.

Our strategy addresses the following complication: we anticipate that the PSF may be distorted from the idealized shape assumed especially with increasing amounts of glycerol [38]. However, the same (possibly incorrect) PSF is used in both FCS and our method in order to compare both methods head-to-head. Thus, concretely, we are asking: how many photon arrivals do we need to converge to the *same* result as FCS (irrespective of whether the FCS result is affected by PSF distortion artifacts)?

Our single photon traces are obtained under a range of conditions, namely different: i) dye concentrations, Fig. (8); ii) diffusion coefficients, Fig. (9); and iii) laser powers, Fig. (10). As before, longer traces, higher concentrations, lower diffusion coefficients, and higher laser powers result, on average, in sharper estimates with the results still converging with at least 2 orders of magnitude fewer photon arrivals than FCS for equal accuracy in Figs. (8), (9), and Fig. (10), respectively. We mention “on average” as individual traces are stochastic. Thus, some traces under higher concentrations of fluorescent molecules may happen to have fewer molecules contribute photons to the traces than experiments with lower concentrations.

Figures (8) recapitulates our expectations derived from the synthetic data shown earlier (Fig. (5)), where dye concentrations are low yielding a wider posterior for our diffusion coefficient and correspondingly sharper posteriors for the higher concentration. Here, similarly to Fig. (1), we compare our method’s diffusion coefficient estimate to FCS as a function of the number of photon arrivals used in the analysis, Fig. (8A) and Fig. (8B), both in good agreement with FCS estimates, produced by the entire traces which is ≈ 10^3^ times longer.

Similar to the analysis of synthetic data, by comparing different diffusion coefficients, the slower a diffusing molecule is, the more time it spends within the confocal volume, the more photons are collected providing us with a sharper posterior estimate of its diffusion coefficient (see Fig. (9)).

Similarly to the synthetic data shown earlier (Fig. (7)), Fig. (10) illustrates the robustness of our method to lower laser power which, as expected, yields a wider posterior for our diffusion coefficient and correspondingly sharper posteriors for higher laser power. Here, we compare our method’s diffusion coefficient estimate to FCS as a function of the number of photon arrivals used in the analysis, Fig. (10A) and Fig. (10B), both in good agreement with FCS estimates, produced from the entire trace.

As further controls, Fig. (11) demonstrates a set of analysis where the background photon emission rate is artificially added to real data. In these cases, we test the limits of our method on more challenging imaging conditons. Furthermore, we repeat our analysis on single photon traces produced by a labeled biomolecule. Specifically, in Fig. (12), we use streptavidin proteins labeled with Cy3.

#### 2. Benchmarking on Experimental Data using 5-TAMRA

Finally, we switch to a different dye, different setup and acquisition electronics as detailed in the Methods section. Our sample contained 20 nM of 5-TAMRA dissolved in water. As previously, we successfully benchmark our estimates of the diffusion coefficient versus the value obtained from FCS on much longer (≈ 10 min) traces, see Fig. (13).

## IV. DISCUSSION

A single photon arriving at a detector mounted to a confocal microsocope encodes information that reports on the fastest timescale achievable for spectroscopic and imaging applications [59, 73]. Directly exploiting this information can help uncover the dynamics of physical or biological systems at fast timescales with accuracy superior to that obtained from *derived* quantities such as down-sampled intensity traces.

Our method takes a Bayesian nonparametrics (BNPs) approach to tackling single photon arrival data to characterize dynamical quantities from as few as hundreds to thousands of datapoints from confocal imaging. This is by contrast to conventional autocorrelative methods used in FCS [30, 69, 84, 104] that require dramatically more data, *i.e.*, datasets several orders of magnitude larger in either total duration or total number of photon arrivals, to characterize dynamical quantities with similar accuracy.

There have been partial solutions to the challenge of interpreting single molecule data at the single photon level often outside FCS applications. Indeed, existing methods make assumptions that render them inapplicable to diffusion through inhomogenesouly illuminated volumes. For example, they assume uniform illumination [40, 82], apply downsampling or binning and thereby reduce temporal resolution to exploit existing mathematical frameworks such as the hidden Markov model [2, 17, 41, 76, 106], or focus on immobile molecules [26, 27, 39, 41]. More recently, fluorescence-based nanosecond FCS approaches, in which the data are still correlated under the assumption that the time trace reports on processes at equilibrium, have been used to obtain information on rapid fluctuations in proteins [90]. As such, correlative methods largely continue to dominate confocal data analysis almost half a century beyond their inception [30, 57, 69].

To take full advantage of single photon data, new Mathematics are required. These must treat the inherent non-stationarity between photon arrivals arising due to molecular diffusion in an inhomogeneously illuminated volume and the stochastic number of molecules contributing photons. In particular, analyzing data derived from mobile molecules within an illuminated confocal region breaks down the perennial parametric Bayesian paradigm that has been the workhorse of data analysis [17, 46, 60, 65, 76, 100, 105]. We argue here that BNPs–which provide principled extensions of the Bayesian methodology [34, 102]–show promise in Physics [48, 54, 60, 95, 96, 98, 100] and give us a working solution to fundamental parametric challenges.

Our new tools open up the possibility to explore at the single photon level non-equilibrium processes resolved on fast timescales [3, 74], reaching *ms* or even below, that have been the focus of recent attention [25]. Moreover, and of immediate relevance for biophysical applications, if a single molecule photobleaches after emitting just a few hundred photons, then our novel method can still provide a diffusion coefficient estimate. Additionally, by analyzing single photon data pointwise, as we do in this study, we obtain a better handle on error bars than analyzing post-processed, such as correlated, data where the error bars can become difficult to compute or interpret [56, 87]. As such, a sharp diffusion coefficient posterior may not only suggest a good estimate of the diffusion coefficient but also suggest that the underlying model, such as normal diffusion, is appropriate and *vice versa* a broad posterior may suggest a poor estimate or an inappropriate motion model.

Furthermore, armed with a transformative framework, founded upon rigorous Statistics, it is now possible to extend the proof-of-principle study to treat effects that lie beyond the current scope of this work. In particular, we can extend our framework to treat multiple color imaging [24], triplet effect and complex molecule photophysics [43] (such as molecular blinking [97, 113] and photobleaching [61, 99]), more complex molecule motion models [101, 112] other than free diffusion [53], distorted or abberated PSF models [32], or even incorporate chemical reactions among the molecules [10, 109]. As our BNP framework explicitly represents the instantaneous position of each involved molecule throughout the experiment’s time course, these are extensions that require modest modifications.

## ACKNOWLEDGEMENTS

S.P. acknowledges support from the NIH NIGMS (R01GM130745) for supporting early efforts in nonparametrics and NIH NIGMS (R01GM134426) for supporting single photon efforts. S.S. acknowledges support from the NIH NIGMS (R01GM121885).

## AUTHOR CONTRIBUTIONS

MT analyzed data and developed analysis software; MT, SJ, IS developed computational tools; OS, SS, BD, ML contributed experimental data; MT, SJ, IS, SP conceived research; SP oversaw all aspects of the projects.

**FIG. A1.**
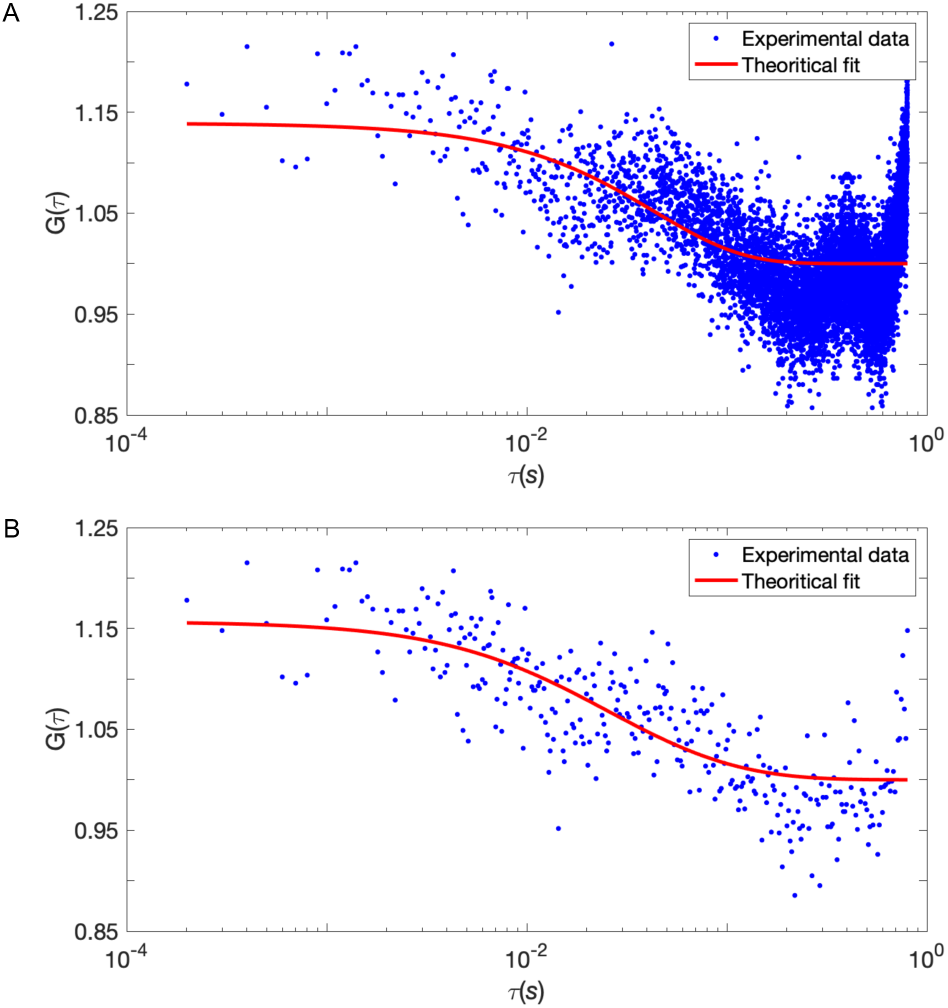
FCS curves resulting from exceedingly short traces (same synthetic data as Fig. 1) with linear (A) and semi-logarithmic (B) binning. Due to the limited data, the quality of the fitted autocorrelation curve, *G*(*τ*), does not improve considerably for (B) as compared to (A).

## Appendix

### Additional Analysis Results

In Fig. (A2) we illustrate the weakness of FCS analysis when applied on limited datasets such as those in the scope of our method. Additionally, using synthetic data, in Fig. (A3) we estimate diffusion coefficients faster than those in the Results section and in Fig. (A4) we estimate photon emission rates. Finally, using experimental data of Cy3 and 5-TAMRA dyes obtained as described in the Methods section, in Figs. (A5) and (A6), respectively, we benchmark the same estimates on real data.

**FIG. A2.**
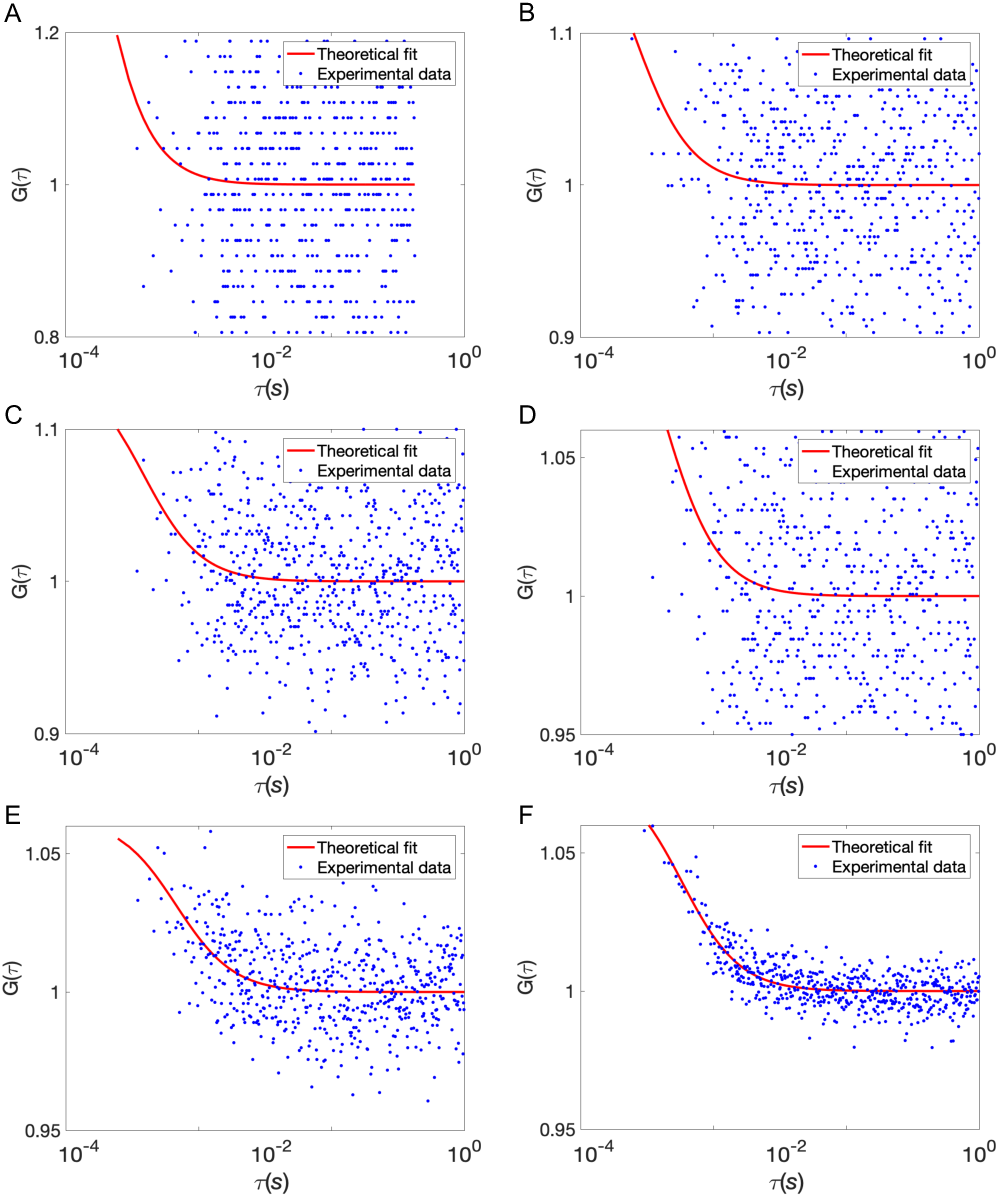
FCS curves resulting from exceedingly short traces. Shown are autocorrelation curves, *G*(*τ*), of 5-TAMRA experimental traces, binned at 10 *μs*, for 100 ms and ≈ 500 photon arrivals (A); 200 ms and ≈ 1000 photon arrivals (B); 300 ms and ≈ 3000 photon arrivals (C); 2 s and ≈ 15000 photon arrivals (D); 30 s and ≈ 15 × 10^5^ photon arrivals (E); 100 s and ≈ 15 × 10^6^ photon arrivals (F). Even a visual inspection illustrates how poorly FCS applies on traces as sort as those analyzed by our BNP method.

**FIG. A3.**
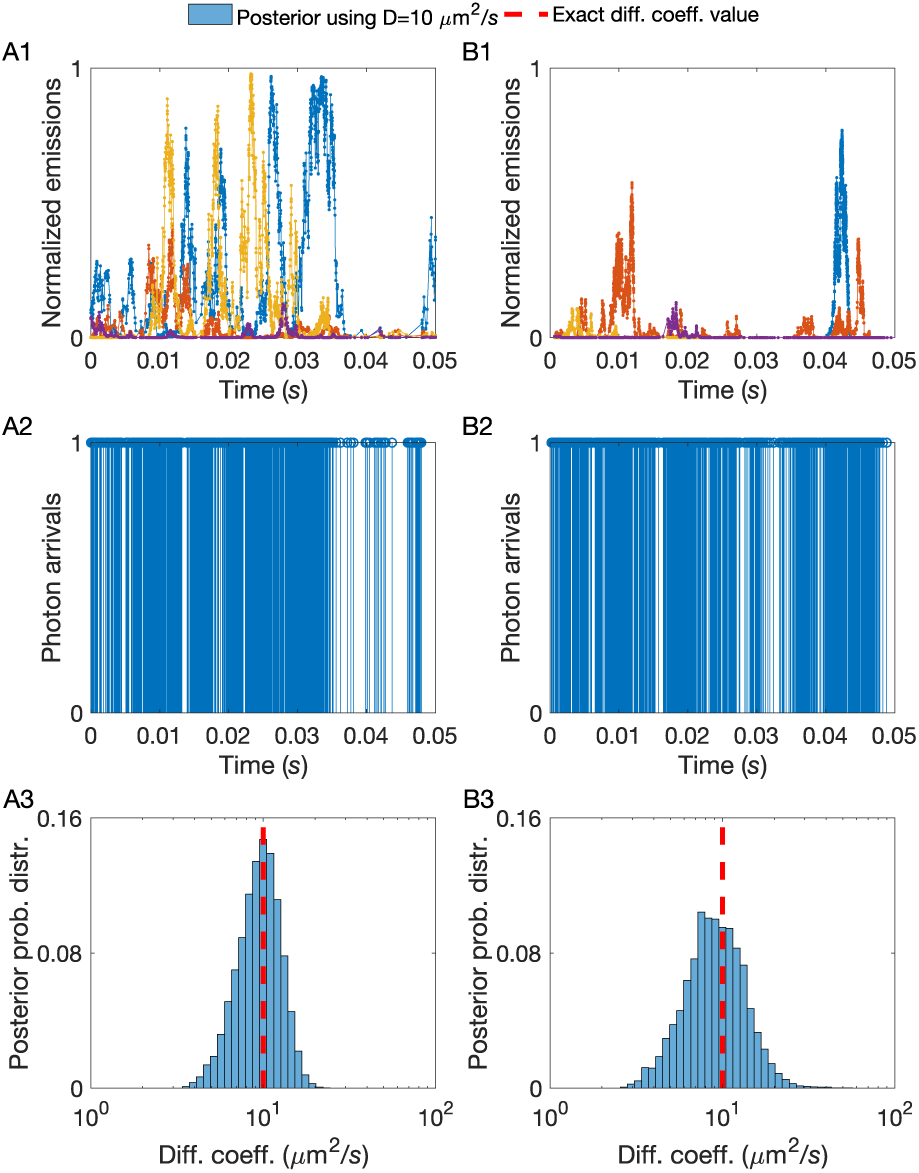
A larger molecule photon emission rate provides more photons per unit time and sharper diffusion coefficient estimates. (A1, B1) Instantaneous molecule photon emission rates 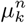, normalized by *μ*_*mol*_. (A2, B2) Photon arrival trace resulting from combining photon emissions from every molecule and the background. These traces are produced by 10 molecules diffusing at 10 *μm*^2^*/s* for a total time of 50 *ms* under background photon emission rate of 10^3^ photons/*s* and molecule photon emission rate 4 × 10^5^ photons*/s* containing ≈ 3000 photon arrivals (A2), and molecule photon emission rate 4 × 10^4^ photons*/s* containing ≈ 2000 photon arrivals (B2). (A3, B3) Posterior probability distributions drawn from traces with differing molecule photon emission rates (shown in (A2, B2)). As expected, for the traces with higher molecule photon emission rate, the peak of the posterior sharply matches with the exact value of *D* (dashed line). Gradually, as we decrease the molecule photon emission rate, the estimation becomes less reliable.

**FIG. A4.**
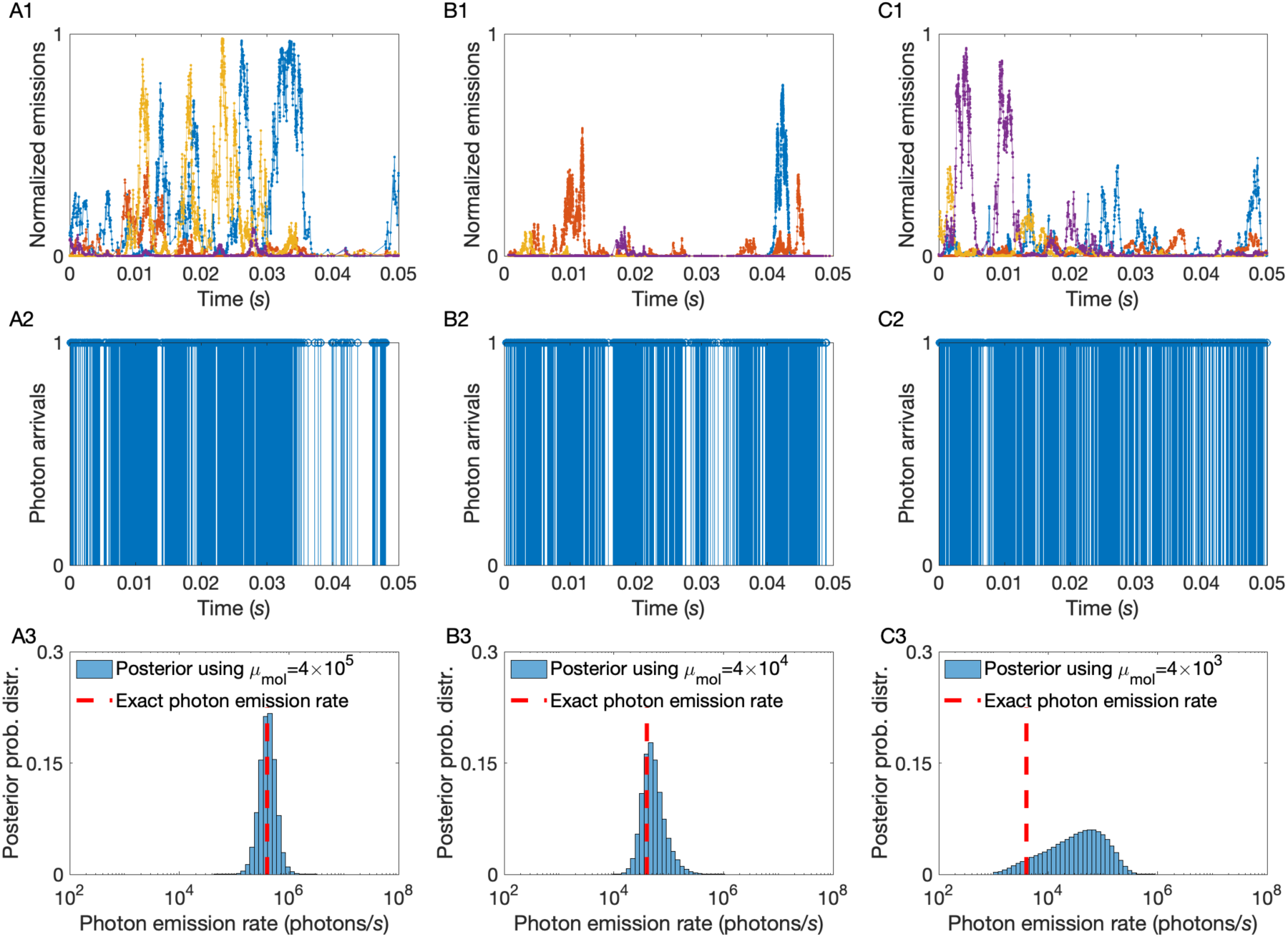
A higher molecule photon emission rate provides more photons per unit time and sharper emission rate estimates. (A1, B1, C1) Instantaneous molecule photon emission rates 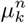, normalized by *μ*_*mol*_. (A2, B2, C2) Photon arrival traces resulting from combining photon emissions from every molecule and the background. These traces produced by 10 molecules diffusing at 10 *μm*^2^*/s* for a total time of 50 *ms* under background photon emission rate of 10^3^ photons/*s* and molecule photon emission rate 4 × 10^5^ photons*/s* containing ≈ 3000 photon arrivals (A2), molecule photon emission rate 4 × 10^4^ photons*/s* containing ≈ 2000 photon arrivals (B2), and molecule photon emission rate 4 × 10^3^ photons*/s* containing ≈ 1000 photon arrivals (C2). (A3, B3, C3) Posterior probability distributions drawn from traces with differing molecule photon emission rates (shown in (A2, B2, C2)). As expected, for the traces with higher molecule photon emission rate, the peak of the posterior sharply matches with the exact value of *μ*_*mol*_ (dashed line). Gradually, as we decrease the molecule photon emission rate, the estimation becomes less reliable.

**FIG. A5.**
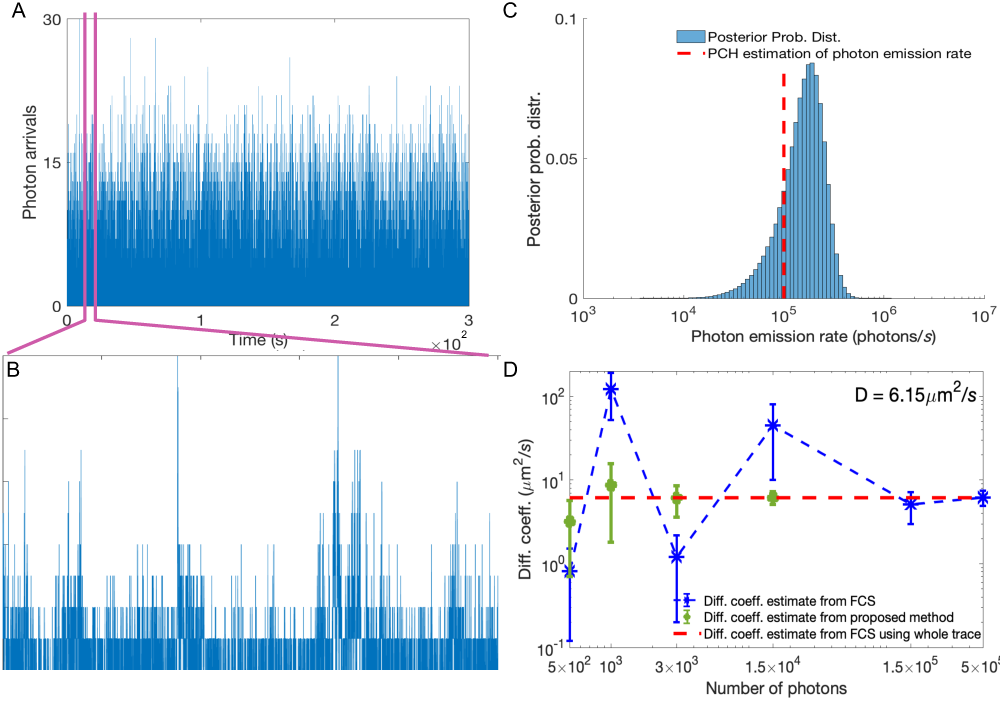
Estimation of the diffusion coefficient and molecule photon emission rate for Cy3 dyes. (A) Experimental intensity trace (binned at 100 *μ*s) with concentration 1 nM of Cy3 dye molecules and 61% glycerol. A background photon emission rate of 600 photons*/s* is known from calibration. (B) Analyzed portion of the trace containing ≈ 3000 photon arrivals. (C) Posterior probability distributions and the value (red dash line) of molecule photon emission rate determined by the photon counting histogram (PCH) method on the entire trace [22]. (D) Similarly to Fig. (1), we compare our method’s diffusion coefficient estimate (green dots) to FCS (blue asterisk) as a function of the number of photons used in the analysis. Since by 1.5 × 10^4^ photon arrivals our method has converged, we avoid analyzing larger traces. The red dash line is the FCS estimate obtained from the entire, 5 min, trace containing ≈ 3 × 10^6^ photon arrivals.

**FIG. A6.**
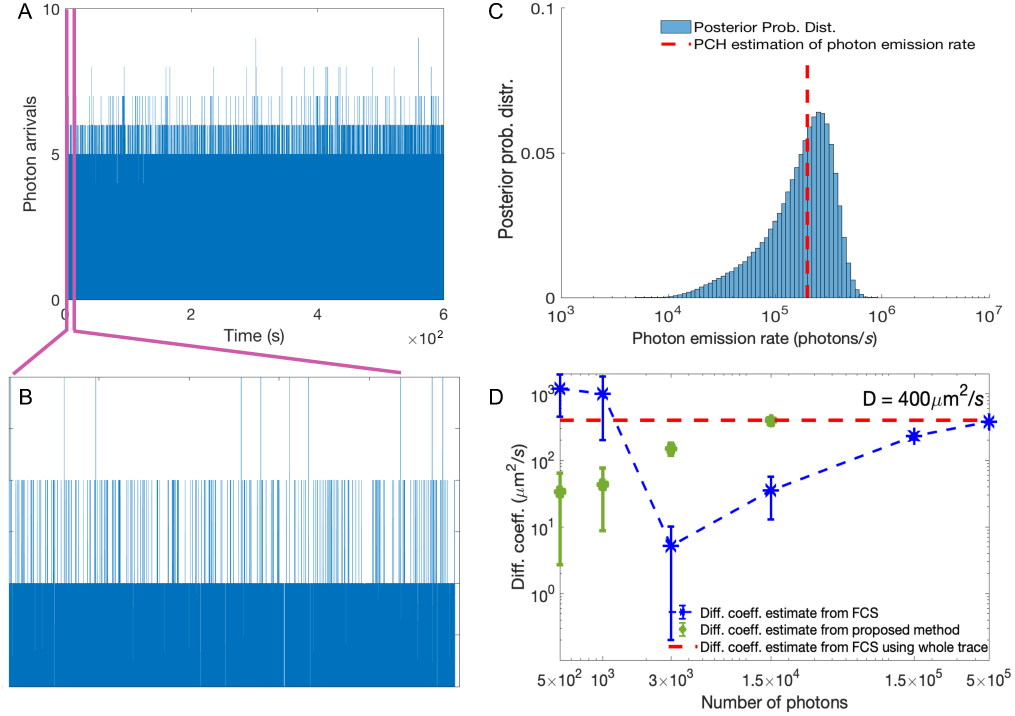
Estimation of the diffusion coefficient and molecule photon emission rate for 5-TAMRA dyes. (A) Experimental intensity trace (binned at 10 *μ*s) with concentration 20 nM of 5-TAMRA dye molecules. A background photon emission rate of 300 photons*/s* is known from calibration. (B) Analyzed portion of the trace containing ≈ 8000 photon arrivals. (C) Posterior probability distributions and the value (red dash line) of molecule photon emission rate determined by the photon counting histogram (PCH) method on the entire trace [22]. (D) Similarly to Fig. (1), we compare our method’s diffusion coefficient estimate (green dots) to FCS (blue asterisk) as a function of the number of photons used in the analysis. Since by 1.5 × 10^4^ photon arrivals our method has converged, we avoid analyzing larger traces. The red dash line is the FCS estimate obtained from the entire, 10 min, trace containing ≈ 6 × 10^6^ photon arrivals.

### Detailed Methods Description

#### Description of Fluorescence Correlation Spectroscopy (FCS)

In FCS the primary quantity of interest is the spontaneously fluctuating fluorescence intensity [29, 91]. Correlations in fluorescence intensities are used to determine physical parameters such as diffusion coefficients. The normalized time autocorrelation function of the fluorescence intensity is defined as

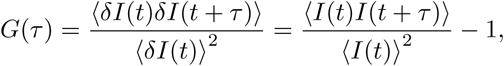

where *I*(*t*) is the fluorescence intensity, *δI*(*t*) is intensity fluctuations at time *t*, and *τ* is the lag time. The intensity fluctuations of the fluorescence intensity are defined as the deviations from the average of the intensity, *δI*(*t*) = *I*(*t*) − ⟨*I*(*t*)⟩. For freely diffusing molecules in a 3D Gaussian confocal volume, the autocorrelation, which we use in this study, is

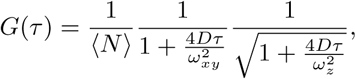

where ⟨*N*⟩ is the average number of molecules in the confocal volume, *D* is the diffusion coefficient, *ω*_*xy*_ and *ω*_*z*_ are the confocal volume axes along the *xy* and *z* directions. Further details on correlative analysis are contained in the cited literature [15, 29, 30, 35, 69, 91].

#### Explanation of Data Simulation

To generate synthetic traces we simulate molecules moving through a three dimensional illuminated volume. The number of moving molecules, *N*, is predefined in each simulation. We apply periodic boundaries to our volume of *L*_*xy*_ and *L*_*z*_ parallel to the focal plane and optical axis, respectively, to keep a relatively stable concentration of molecules near the confocal volume.

We denote the locations of the molecules as 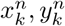 and 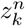, where *k* labels time levels and *n* = 1, 2, …, *N* labels molecules. The total trace duration *T*_*total*_ = *t*_*K*_ − *t*_0_, is predefined. The time intervals between successive recorded photons Δ*t*_*k*−1_ = *t*_*k*_ − *t*_*k*−1_, are generated through pseudo-random computer simulations and recorded for subsequent analysis.

The locations of the molecules 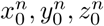 at the first evaluation time *t*_0_ are randomly sampled from the uniform distribution with borders identical to the boundaries ±*L*_*xy*_ and ±*L*_*z*_ of the prescribed simulation region. Locations 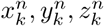, for *k* = 1, …, *K*, at times *t*_*k*_ are generated according to the diffusion model explained above under a predefined diffusion coefficient *D*.

We obtain photon inter-arrival times, **Δ*t*** = (Δ*t*_1_, Δ*t*_2_, …, Δ*t*_*K*−1_), by simulating exponential random variables of rate *μ*_*k*_. For independent background and molecule photon emission rates, the corresponding exponential emission mean rates *μ*_*k*_ depend on a Gaussian PSF as eqs. (2)–(3). Both background, *μ*_*back*_, and the molecule photon emission rate, *μ*_*mol*_, are predefined.

#### Definition of Molecule Photon Emission Rate

In this study the emission rate of detected photons for a single fluorophore at position *x, y, z* is used. This is formulated as the product *μ*(*x, y, z*) = *μ*_0_*φ*_*d*_*φ*_*de*_*φ*_*f*_ *σ*EXC(*x, y, z*)CEF(*x, y, z*). Here, *μ*_0_ and *φ*_*d*_ are the maximum excitation intensity and the efficiency of the photon collection at the center of the confocal volume, respectively, *φ*_*de*_ is the efficiency of the detector, *φ*_*f*_ is the quantum efficiency of the fluorophore, *σ* is the fluorophore absorption cross-section, EXC(*x, y, z*) is the excitation profile and CEF(*x, y, z*) is the detection profile [31]. By revising the definition of *μ*(*x, y, z*), we obtain *μ*(*x, y, z*) = *μ*_*mol*_PSF(*x, y, z*) where *μ*_*mol*_ = *μ*_0_*φ*_*d*_*φ*_*de*_*φ*_*f*_ *σ* and PSF(*x, y, z*) = EXC(*x, y, z*)CEF(*x, y, z*).

To relate our *single molecule* photon emission rate *μ*_*mol*_ to the average photon count rate typically determined in *bulk* experiments, we compute a spatial average

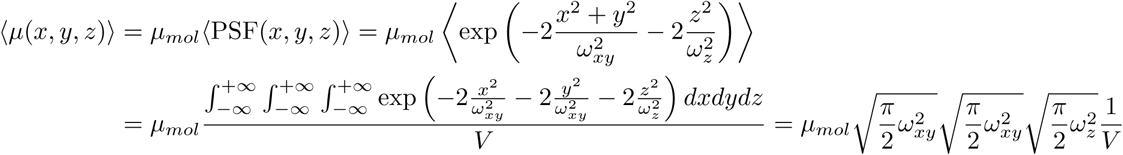

where *V* denotes our PSF’s effective volume [59, 85] which is equal to 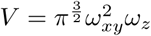. As a result, our molecule photon emission rate *μ*_*mol*_ is related to ⟨*μ*(*x, y, z*)⟩ according to 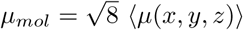.

#### Description of Wilson-Hiferty Approximation

To perform the necessary computations of the next section, we use a Wilson-Hiferty transform [110] to approximate the probability density of exponential random variables. We use this approximation to sample the locations of the molecules within our overall Gibbs sampler (see next).

To apply the Wilson-Hiferty approximation, first we transform our observation random variable Δ*t*_*k*_ to a new random variable *ρ*_*k*_, where *ρ*_*k*_ = 2*μ*_*k*_Δ*t*_*k*_. A change of variables, indicates that Δ*t*_*k*_|*μ*_*k*_ ∼ Exponential(*μ*_*k*_) implies *ρ*_*k*_|*μ*_*k*_ ∼ *χ*^2^ (2), where *χ*^2^(2) denotes the chi-square probability distribution with 2 degrees of freedom. By applying another transformation, where 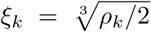, according to [110], *ξ*_*k*_ follows an *approximately* normal probability distribution 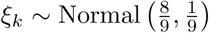. So, by 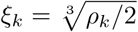 and *ρ*_*k*_ = 2*μ*_*k*_Δ*t*_*k*_, we conclude 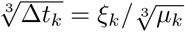. Therefore, since 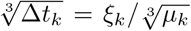, we establish the approximation

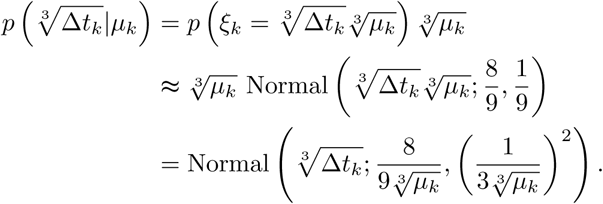

### Detailed Description of the Inference Framework

#### Prior Probability Distributions

Within the Bayesian paradigm, all unknown model parameters need priors. These parameters are: the diffusion coefficient *D*; the molecule photon emission rate *μ*_*mol*_; the initial molecule locations 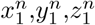; as well as the indicator prior weights *q*^*n*^.

##### Prior on the Diffusion Coefficient

To make sure that *D* sampled in our formulation attains only positive values, we choose an Inverse-Gamma prior

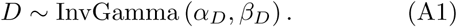

This prior is conjugate to the motion model which simplifies the computations shown below.

##### Priors on Molecule Photon Emission Rate

To guarantee that *μ*_*mol*_ sampled in our formulation also attains only positive values, we choose a Gamma prior

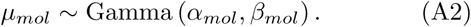

##### Priors on Initial Molecule Locations

Because of the symmetries inherent to the confocal volume, *e.g.*, a molecule at a location (*x, y, z*) gives rise to the same photon emission rate as a molecule at location, (−*x*, −*y*, −*z*), we use priors on the initial locations that respect these symmetries. To simplify the computations, we use independent symmetric normal distributions

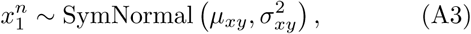

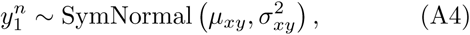

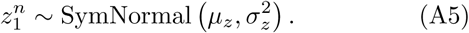

##### Priors and Hyperpriors for the Indicators

To simplify the computations described in the next section, we use a finite, but large, model population of *N* molecules that contain contributing and noncontributing ones. These molecules are collectively indexed by *n* = 1, 2, …, *N*. As described in the Methods section, inferring how many molecules are actually warranted by the data analyzed is the same as estimating how many of those *N* molecules are active, *i.e., b*^*n*^ = 1, while the rest are inactive, *i.e., b*^*n*^ = 0, and so have no influence and are applied just for computational reasons.

We use a Bernoulli prior of weight *q*^*n*^ to make sure that each indicator *b*^*n*^ takes only values 0 or 1. Moreover, on each weight *q*^*n*^, we assign a beta hyperprior

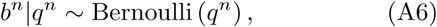

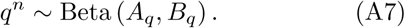

To make sure that the resulting formulation avoids overfitting, we make the specific selections *A*_*q*_ = *α*_*q*_*/N* and *B*_*q*_ = *β*_*q*_(*N* − 1)*/N*. For these choices [1, 16, 77, 78], and in the limit that *N* → ∞ (that is, when the assumed molecule population is large), this prior/hyperprior choice converges to a non-parametric beta-Bernoulli process. Therefore, for *N* ≫ 1, the posterior is well defined and becomes *independent* of the selected value of *N*. In other words, provided *N* is large enough, its effect on the results is negligible; while its precise value has only computational implications.

#### Description of the Computational Implementation

Here, 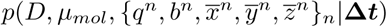 is the joint probability distribution of our framework where molecular trajectories and measurements are gathered in

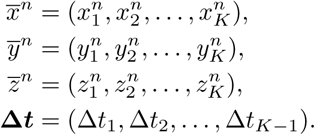

Posterior samples are generated according to Gibbs sampling [37, 66, 86, 100, 107]. We achieve this by sampling a variable conditioned on all other variables and the given photon inter-arrival times **Δ*t***. Conceptually, the steps in the generation of each posterior sample 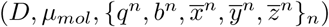 are:

1. For each *n* of the *active* molecules
  a. Update trajectory 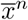 of active molecule *n*
  b. Update trajectory 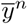 of active molecule n
  c. Update trajectory 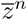 of active molecule *n*
2. Update jointly the trajectories 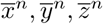 for all n of the inactive molecules
3. Update the diffusion coefficient D
4. Update jointly the prior weights q^n^ for all model molecules and simultaneously update jointly the indicators b^n^ for all model molecules
5. Update the molecule photon emission rate *μ*_*mol*_

##### Sampling Active Molecules Locations

To sample the location of an active molecule 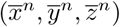, we use forward filtering and backward sampling [11, 20, 53, 94]. In particular, we update each dimension sequentially from the following full conditional probability distributions 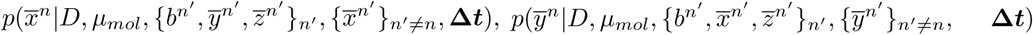,, and 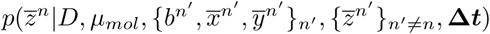. Below, we show in detail the calculation only for sampling 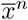, since for sampling 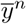 and 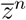 they are similar.

To sample the trajectory 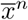, we rely on the factorization

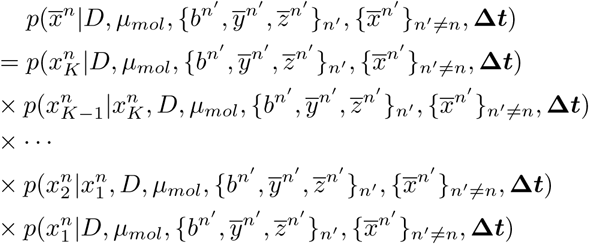

and, according to this factorization, we sample individual locations 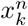 sequentially

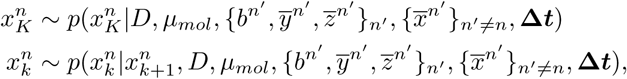

where, *k* = 1, …, *K* − 1. However, to be able to perform these steps, we first need to compute the involved probability distributions. We describe below a computationally efficient way to do so that proceeds in a forward filtering and a backward sampling step.

Before we start the sampling of the locations, we determine each one of the individual probability distributions that are needed. To do this in a computationally tractable manner [14, 20], we compute filter distributions 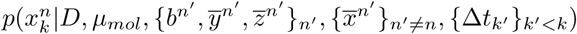.

In our case, both dynamic (eqs. (7)–(9))) and observation (eq. (1)) probability distributions provide equal probabilities for 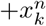 and 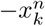. Therefore, the filter distribution consists of two modes symmetrically placed across the origin [53]. Accordingly, we compute an *approximate* bimodal symmetric filter of the form

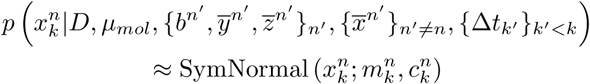

where SymNormal 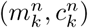 describes the symmetric normal distribution. The filter, that is the values of 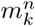 and 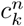, is updated iteratively according to

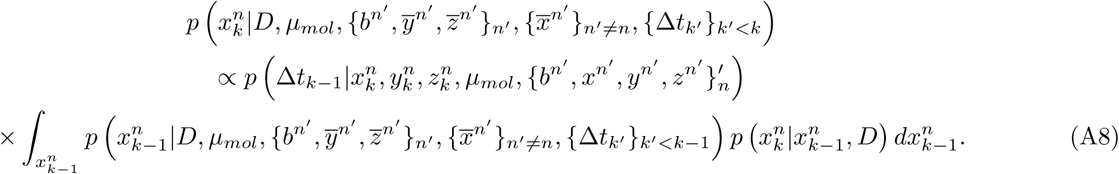

To be able to carry out these computations efficiently, similar to [53], we work on an approximate model where the exponential emission equation, eq. (1), is replaced by a normal one using the Wilson-Hiferty approximation as we discussed earlier. Our *approximate* emission equation is

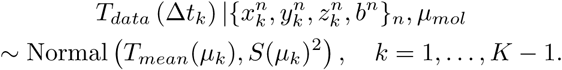

where *μ*_*k*_ is given in eq. (2); while *T*_*data*_ (Δ*t*_*k*_), *T*_*mean*_(*μ*_*k*_) and *S*^2^(*μ*_*k*_) are given by the Wilson-Hiferty approximation [110]. As explained earlier, the approximation is given by 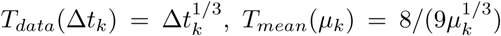 and 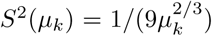. Because of the specific choices of our problem (*i.e.*, diffusive molecules, symmetric normal filter at the proceeding time, and normal likelihood), eq. (A8) reduces to

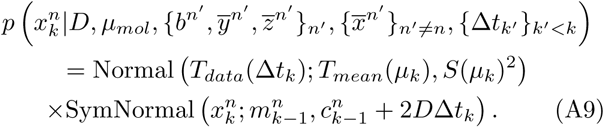

Finally, to obtain the values of 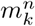 and 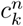, we linearize the product in eq. (A9) as described next. From eq. (A9), we have

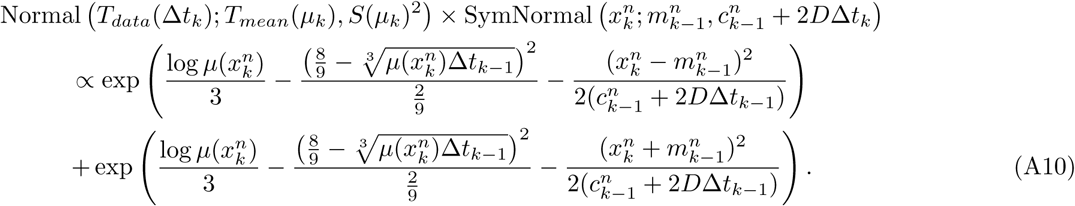

The density in eq. (A10) consists of two modes, one on the positive semi-axis of *x* and one on the negative semi-axis of *x*. Considering 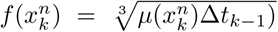 and 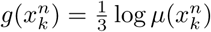 and linearizing them around the previous filter’s mode, 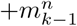 or 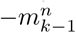, the modes of eq. (A10) are approximated by

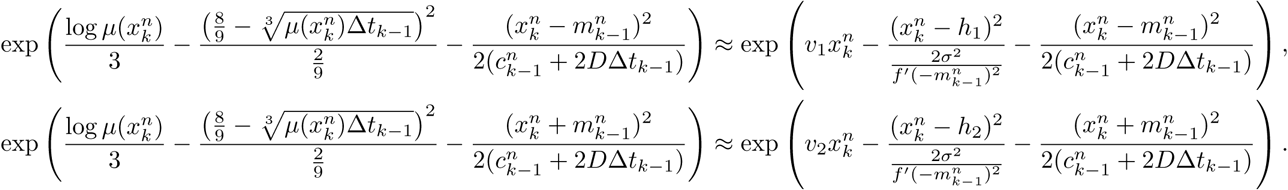

Combining both approximations, the density of eq. (A10), is approximated by

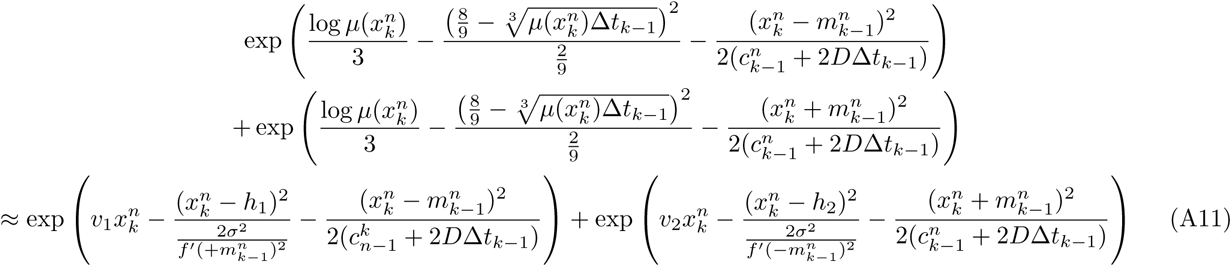

where 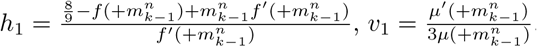 and 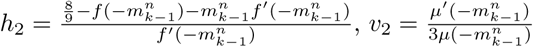.

Equation (A11) describes a symmetric normal distribution. Equating this distribution with our filter, *i.e.*, SymNormal 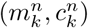, we obtain 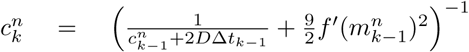, and 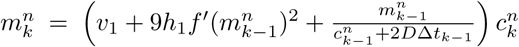. These apply for *k* = 2, …, *K* and are used to update the filter. To begin, we use eq. (A3)–(A5) and set 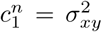 and 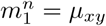.

Having computed the filter distributions above, we are able to sample the individual locations by starting from 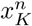 and moving backward towards 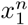. In particular, by applying the Bayes’ rule, each one of the individual distributions factorize as

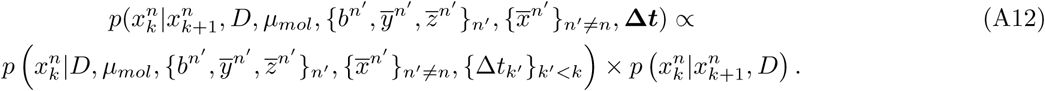

The first term is given by the filter distribution which is replaced by our approximate SymNormal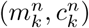, and the second term is our motion model Normal 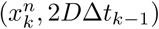, all of which are known at this stage. Therefore, backward sampling starts at 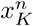 and continues for 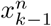 with

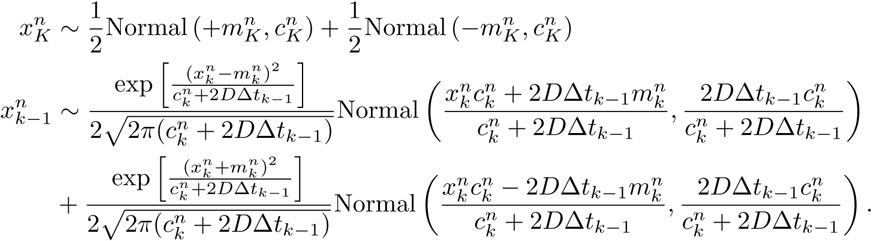

##### Sampling Inactive Molecule Trajectories

To update the trajectories of the inactive molecules, we sample from the corresponding conditionals 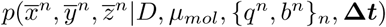. Since the inactive molecules are not associated with the observations in **Δ*t***, these reduce to 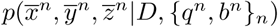 which we simulate as standard 3D Brownian motion [64].

##### Sampling the Diffusion Coefficient

We sample the diffusion coefficient from the conditional probability distribution 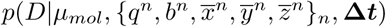. Because of the specific dependencies of the variables in this formulation, *e.g.*, eq. (A1) and eqs. (7)–(9), the conditional distribution simplifies to 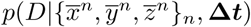. Using Bayes’ rule, this distribution becomes 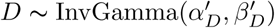 where 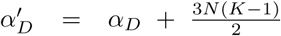 and 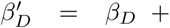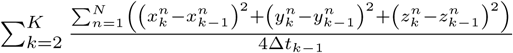.

##### Sampling Molecule Indicators

For each molecule *n* we sample its indicator prior weight, *q*^*n*^, from the corresponding conditional distribution 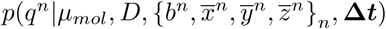, which simplifies to *p*(*q*^*n*^|*b*^*n*^). For this we use eq. (A7) and eq. (A6). According to Bayes’ rule, the latter distribution becomes *q*^*n*^ ∼ Beta(*α*′, *β*′) where 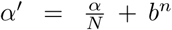 and 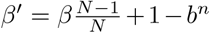. Subsequently, we update the indicators *b*^*n*^ by sampling from the corresponding conditional distribution 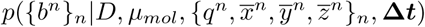 using a Methropolis-Hasting algorithm [18, 23]. For this, we use a proposal 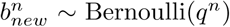. With this proposal, the acceptance ratio becomes

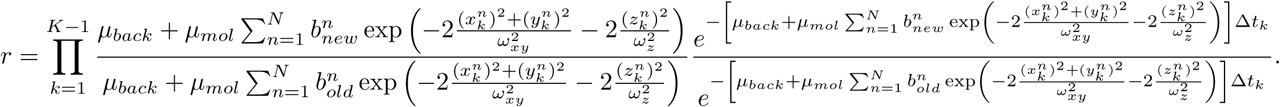

##### Sampling the Molecule Photon Emission Rate

In the last step, after updating the locations and indicators of the molecules, we sample the molecule photon emission rate from the corresponding conditional distribution 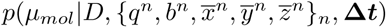. To sample this distribution, we also use a Metropolis-Hastings step. For this, we use proposals of the form 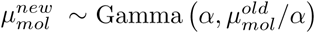 where 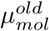 denotes the current sampled value. Using both eqs. (1) and (2), the acceptance ratio becomes

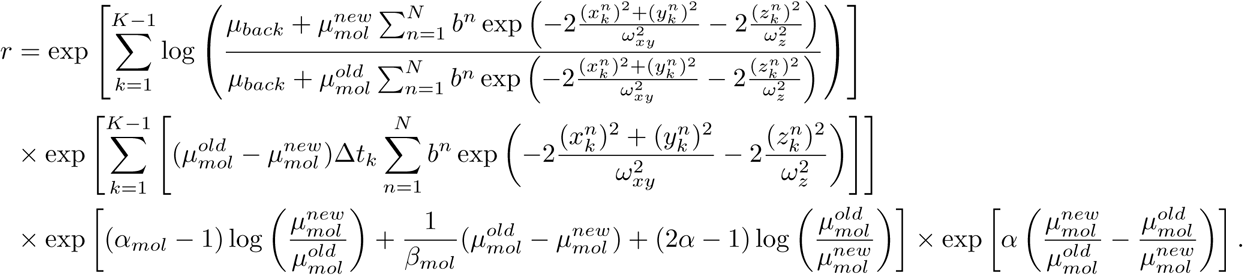

---

**TABLE II.** Summary of notation.

| Description | Variable | Units |
| --- | --- | --- |
| Diffusion coefficient | $D$ | $\mu\text{m}^2/\text{s}$ |
| $\alpha$ parameter of the diffusion coefficient prior | $\alpha_D$ | - |
| $\beta$ parameter of the diffusion coefficient prior | $\beta_D$ | $\mu\text{m}^2/\text{s}$ |
| Photon inter-arrival time | $\Delta t$ | s |
| Total trace duration | $T_{total}$ | s |
| molecule photon emission rate (maximum) | $\mu_{mol}$ | photons/s |
| $\alpha$ parameter of the molecule photon emission rate's prior | $\alpha_{mol}$ | - |
| $\beta$ parameter of the molecule photon emission rate's prior | $\beta_{mol}$ | photons/s |
| Emission rate of molecule $n$ at time $t_k$ | $\mu_k^n$ | photons/s |
| Combined photon emission rate at time $t_k$ | $\mu_k$ | photons/s |
| Background photon emission rate | $\mu_{back}$ | photons/s |
| Minor semi-axis of confocal PSF (focal plane) | $\omega_{xy}$ | $\mu\text{m}$ |
| Major semi-axis of confocal PSF (optical axis) | $\omega_z$ | $\mu\text{m}$ |
| Location of molecule $n$ at time $t_k$ in $x$-coordinate | $x_k^n$ | $\mu\text{m}$ |
| Location of molecule $n$ at time $t_k$ in $y$-coordinate | $y_k^n$ | $\mu\text{m}$ |
| Location of molecule $n$ at time $t_k$ in $z$-coordinate | $z_k^n$ | $\mu\text{m}$ |
| Recorded photon inter-arrival time between $t_k$ and $t_{k-1}$ | $\Delta t_k$ | s |
| Indicator variable for molecule $n$ | $b^n$ | - |
| Prior weight for $b_n$ | $q^n$ | - |
| $\alpha$ parameter of prior weight $q^n$ | $\alpha_q$ | - |
| $\beta$ parameter of prior weight $q^n$ | $\beta_q$ | - |
| Upper bound for the number of model molecules | $N$ | - |
| Mean value of initial molecule position's prior in the $xy$-plane | $\mu_{xy}$ | $\mu\text{m}$ |
| Mean value of initial molecule position's prior on the $z$-axis | $\mu_z$ | $\mu\text{m}$ |
| Variance of the initial molecule position's prior in the $xy$-plane | $\sigma_{xy}^2$ | $\mu\text{m}$ |
| Variance of the initial molecule position's prior on the $z$-axis | $\sigma_z^2$ | $\mu\text{m}$ |
| Periodic boundary in the $xy$-plane | $L_{xy}$ | $\mu\text{m}$ |
| Periodic boundary on the $z$-axis | $L_z$ | $\mu\text{m}$ |

**TABLE III.** Probability distributions used and their densities. Here, the corresponding random variables are denoted by *x*. We use “;” to separate random variables from parameters. For example, Normal(*x*; *μ, σ*^2^) means that *x* is the random variable (e.g. 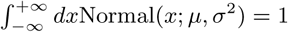), and *μ* and *σ*^2^ are parameters characterizing this density.

| Distribution | Notation | Probability density function | Mean | Variance |
| --- | --- | --- | --- | --- |
| Normal | $\text{Normal}(\mu, \sigma^2)$ | $\frac{1}{\sqrt{2\pi\sigma^2}} e^{-\frac{(x-\mu)^2}{2\sigma^2}}$ | $\mu$ | $\sigma^2$ |
| Symmetric Normal | $\text{SymNormal}(\mu, \sigma^2)$ | $\frac{1}{2} \frac{e^{-\frac{(x+\mu)^2}{2\sigma^2}}}{\sqrt{2\pi\sigma^2}} + \frac{1}{2} \frac{e^{-\frac{(x-\mu)^2}{2\sigma^2}}}{\sqrt{2\pi\sigma^2}}$ | 0 | $\mu^2 + \sigma^2$ |
| Exponential | $\text{Exponential}(\mu)$ | $\mu e^{-\mu x}$ | $\frac{1}{\mu}$ | $\frac{1}{\mu^2}$ |
| Chi-square | $\chi^2(\alpha, 2)$ | $\frac{1}{\Gamma(\frac{\alpha}{2}) 2^{\frac{\alpha}{2}}} x^{\frac{\alpha}{2}-1} e^{-\frac{x}{2}}$ | $\alpha$ | $2\alpha$ |
| Gamma | $\text{Gamma}(\alpha, \beta)$ | $\frac{1}{\Gamma(\alpha)\beta^\alpha} x^{\alpha-1} e^{-\frac{x}{\beta}}$ | $\alpha\beta$ | $\alpha\beta^2$ |
| Inverse-Gamma | $\text{InvGamma}(\alpha, \beta)$ | $\frac{\beta^\alpha}{\Gamma(\alpha)} x^{-\alpha-1} e^{-\frac{\beta}{x}}$ | $\frac{\beta}{\alpha-1}$ | $\frac{\beta^2}{(\alpha-1)^2(\alpha-2)}$ |
| Beta | $\text{Beta}(\alpha, \beta)$ | $\frac{\Gamma(\alpha+\beta)}{\Gamma(\alpha)\Gamma(\beta)} x^{\alpha-1} (1-x)^{\beta-1}$ | $\frac{\alpha}{\alpha+\beta}$ | $\frac{\alpha\beta}{(\alpha+\beta)^2(\alpha+\beta+1)}$ |
| Bernoulli | $\text{Bernoulli}(q)$ | $(q-1)\delta_0(x) + q\delta_1(x)$ | $q$ | $q(1-q)$ |

**TABLE IV.** Parameter values used in the generation of the synthetic traces. Choices are listed according to figures.

| Units | $L_{xy}$<br>$\mu m$ | $L_z$<br>$\mu m$ | $\omega_{xy}$<br>$\mu m$ | $\omega_z$<br>$\mu m$ | $N$<br>- | $D$<br>$\mu m^2/s$ | $\mu_{mol}$<br>photons/s | $\mu_{back}$<br>photons/s | $T_{total}$<br>s |
| --- | --- | --- | --- | --- | --- | --- | --- | --- | --- |
| Fig. (2A) | 1 | 2 | 0.3 | 1.5 | 4 | 1 | $4 \times 10^4$ | $10^3$ | 0.03 |
| Fig. (2B) | 1 | 2 | 0.3 | 1.5 | 4 | 1 | $4 \times 10^4$ | $10^3$ | 0.03 |
| Fig. (2C) | 1 | 2 | 0.3 | 1.5 | 4 | 1 | $4 \times 10^4$ | $10^3$ | 0.03 |
| Fig. (2D) | 1 | 2 | 0.3 | 1.5 | 4 | 1 | $4 \times 10^4$ | $10^3$ | 0.03 |
| Fig. (2E) | 1 | 2 | 0.3 | 1.5 | 4 | 1 | $4 \times 10^4$ | $10^3$ | 0.03 |
| Fig. (4A) | 1 | 2 | 0.3 | 1.5 | 4 | 1 | $4 \times 10^4$ | $10^3$ | 0.03 |
| Fig. (5A) | 1 | 2 | 0.3 | 1.5 | 10 | 1 | $4 \times 10^4$ | $10^3$ | 0.03 |
| Fig. (5C) | 1 | 2 | 0.3 | 1.5 | 4 | 1 | $4 \times 10^4$ | $10^3$ | 0.03 |
| Fig. (5E) | 1 | 2 | 0.3 | 1.5 | 1 | 1 | $4 \times 10^4$ | $10^3$ | 0.03 |
| Fig. (6) | 1 | 2 | 0.3 | 1.5 | 4 | $10^{-2}$ | $4 \times 10^4$ | $10^3$ | 0.03 |
| Fig. (6) | 1 | 2 | 0.3 | 1.5 | 4 | $10^{-1}$ | $4 \times 10^4$ | $10^3$ | 0.03 |
| Fig. (6) | 1 | 2 | 0.3 | 1.5 | 4 | 1 | $4 \times 10^4$ | $10^3$ | 0.03 |
| Fig. (6) | 1 | 2 | 0.3 | 1.5 | 4 | 10 | $4 \times 10^4$ | $10^3$ | 0.03 |
| Fig. (6) | 1 | 2 | 0.3 | 1.5 | 4 | 100 | $4 \times 10^4$ | $10^3$ | 0.03 |
| Fig. (7A) | 1 | 2 | 0.3 | 1.5 | 4 | 1 | $4 \times 10^5$ | $10^3$ | 0.03 |
| Fig. (7B) | 1 | 2 | 0.3 | 1.5 | 4 | 1 | $4 \times 10^4$ | $10^3$ | 0.03 |
| Fig. (7C) | 1 | 2 | 0.3 | 1.5 | 4 | 1 | $4 \times 10^3$ | $10^3$ | 0.03 |
| Fig. (7D) | 1 | 2 | 0.3 | 1.5 | 4 | 1 | $10^3$ | $10^3$ | 0.03 |
| Fig. (A3A) | 1 | 2 | 0.3 | 1.5 | 10 | 10 | $4 \times 10^5$ | $10^3$ | 0.05 |
| Fig. (A3C) | 1 | 2 | 0.3 | 1.5 | 10 | 10 | $4 \times 10^4$ | $10^3$ | 0.05 |
| Fig. (A4A) | 1 | 2 | 0.3 | 1.5 | 10 | 10 | $4 \times 10^5$ | $10^3$ | 0.05 |
| Fig. (A4C) | 1 | 2 | 0.3 | 1.5 | 10 | 10 | $4 \times 10^4$ | $10^3$ | 0.05 |
| Fig. (A4E) | 1 | 2 | 0.3 | 1.5 | 10 | 10 | $4 \times 10^3$ | $10^3$ | 0.05 |

**TABLE V.** Parameter values used in the analyses of the traces. Choices are listed according to figures.

| Units | $\omega_{xy}$<br>$\mu m$ | $\omega_z$<br>$\mu m$ | $N$<br>- | $\alpha_D$<br>- | $\beta_D$<br>$\mu m^2/s$ | $\alpha_{mol}$<br>- | $\beta_{mol}$<br>phts/s | $\alpha_q$<br>- | $\beta_q$<br>- | $\mu_{xy}$<br>- | $\mu_z$<br>$\mu m$ | $\sigma_{xy}^2$<br>$\mu m$ | $\sigma_z^2$<br>$\mu m^2$ |
| --- | --- | --- | --- | --- | --- | --- | --- | --- | --- | --- | --- | --- | --- |
| Fig. (2A) | 0.3 | 1.5 | - | 1 | 1 | 1 | $10^5$ | - | - | 0.1 | 0.1 | 1 | 1 |
| Fig. (2B) | 0.3 | 1.5 | - | 1 | 1 | 1 | $10^5$ | - | - | 0.1 | 0.1 | 1 | 1 |
| Fig. (2C) | 0.3 | 1.5 | - | 1 | 1 | 1 | $10^5$ | - | - | 0.1 | 0.1 | 1 | 1 |
| Fig. (2D) | 0.3 | 1.5 | - | 1 | 1 | 1 | $10^5$ | - | - | 0.1 | 0.1 | 1 | 1 |
| Fig. (2E) | 0.3 | 1.5 | - | 1 | 1 | 1 | $10^5$ | - | - | 0.1 | 0.1 | 1 | 1 |
| Fig. (4) | 0.3 | 1.5 | 20 | 1 | 1 | 1 | $10^5$ | 1 | 1 | 0.1 | 0.1 | 1 | 1 |
| Fig. (5A3) | 0.3 | 1.5 | 20 | 1 | 1 | 1 | $10^5$ | 1 | 1 | 0.1 | 0.1 | 1 | 1 |
| Fig. (5B3) | 0.3 | 1.5 | 20 | 1 | 1 | 1 | $10^5$ | 1 | 1 | 0.1 | 0.1 | 1 | 1 |
| Fig. (5C3) | 0.3 | 1.5 | 20 | 1 | 1 | 1 | $10^5$ | 1 | 1 | 0.1 | 0.1 | 1 | 1 |
| Fig. (6) | 0.3 | 1.5 | 20 | 1 | 1 | 1 | $10^5$ | 1 | 1 | 0.1 | 0.1 | 1 | 1 |
| Fig. (6) | 0.3 | 1.5 | 20 | 1 | 1 | 1 | $10^5$ | 1 | 1 | 0.1 | 0.1 | 1 | 1 |
| Fig. (6) | 0.3 | 1.5 | 20 | 1 | 1 | 1 | $10^5$ | 1 | 1 | 0.1 | 0.1 | 1 | 1 |
| Fig. (6) | 0.3 | 1.5 | 20 | 1 | 100 | 1 | $10^5$ | 1 | 1 | 0.1 | 0.1 | 1 | 1 |
| Fig. (7A) | 0.3 | 1.5 | 20 | 1 | 1 | 1 | $10^5$ | 1 | 1 | 0.1 | 0.1 | 1 | 1 |
| Fig. (7B) | 0.3 | 1.5 | 20 | 1 | 1 | 1 | $10^5$ | 1 | 1 | 0.1 | 0.1 | 1 | 1 |
| Fig. (7C) | 0.3 | 1.5 | 20 | 1 | 1 | 1 | $10^5$ | 1 | 1 | 0.1 | 0.1 | 1 | 1 |
| Fig. (7D) | 0.3 | 1.5 | 20 | 1 | 1 | 1 | $10^5$ | 1 | 1 | 0.1 | 0.1 | 1 | 1 |
| Fig. (8) | 0.23 | 0.55 | 20 | 1 | 1 | 1 | $10^5$ | 1 | 1 | 0.1 | 0.1 | 1 | 1 |
| Fig. (9) | 0.23 | 0.55 | 20 | 1 | 1 | 1 | $10^5$ | 1 | 1 | 0.1 | 0.1 | 1 | 1 |
| Fig. (10) | 0.23 | 0.55 | 20 | 1 | 1 | 1 | $10^5$ | 1 | 1 | 0.1 | 0.1 | 1 | 1 |
| Fig. (11) | 0.23 | 0.55 | 20 | 1 | 1 | 1 | $10^5$ | 1 | 1 | 0.1 | 0.1 | 1 | 1 |
| Fig. (12) | 0.27 | 4.51 | 20 | 1 | 1 | 1 | $10^5$ | 1 | 1 | 0.1 | 0.1 | 1 | 1 |
| Fig. (13) | 0.22 | 3.90 | 20 | 1 | 100 | 1 | $10^5$ | 1 | 1 | 0.1 | 0.1 | 1 | 1 |
| Fig. (A3) | 0.3 | 1.5 | 20 | 1 | 1 | 1 | $10^5$ | 1 | 1 | 0.1 | 0.1 | 1 | 1 |
| Fig. (A4) | 0.3 | 1.5 | 20 | 1 | 1 | 1 | $10^5$ | 1 | 1 | 0.1 | 0.1 | 1 | 1 |
| Fig. (A5) | 0.23 | 0.55 | 20 | 1 | 1 | 1 | $10^5$ | 1 | 1 | 0.1 | 0.1 | 1 | 1 |
| Fig. (A6) | 0.22 | 3.90 | 20 | 1 | 100 | 1 | $10^5$ | 1 | 1 | 0.1 | 0.1 | 1 | 1 |

